# Liquid spherical shells are a non-equilibrium steady state

**DOI:** 10.1101/2023.01.31.526480

**Authors:** Alexander M. Bergmann, Jonathan Bauermann, Giacomo Bartolucci, Carsten Donau, Michele Stasi, Anna-Lena Holtmannspötter, Frank Jülicher, Christoph A. Weber, Job Boekhoven

**Affiliations:** School of Natural Sciences, Department of Chemistry, Technical University of Munich, Lichtenbergstrasse 4, 85748 Garching, Germany; Max Planck Institute for the Physics of Complex Systems, Nöthnitzer Strasse 38, 01187 Dresden, Germany; Center for Systems Biology Dresden, Pfotenhauerstrasse 108, 01307 Dresden, Germany; Faculty of Mathematics, Natural Sciences, and Materials Engineering: Institute of Physics, University of Augsburg, Universitätsstrasse 1, 86159 Augsburg, Germany

**Author notes:** equal contribution (Alexander M. Bergmann, Jonathan Bauermann, Giacomo Bartolucci, Carsten Donau).

**Keywords:** liquid-liquid phase separation, dissipative structures, chemically fueled droplets

## Abstract

Liquid-liquid phase separation is the process in which two immiscible liquids demix. This spontaneous phenomenon yields spherical droplets that eventually coarsen to one large, stable droplet governed by the principle of minimal free energy. In chemically fueled phase separation, the formation of phase-separating molecules is coupled to a fuel-driven, nonequilibrium reaction cycle. Chemically fueled phase separation yields dissipative structures sustained by a continuous fuel conversion. Such dissipative structures are ubiquitous in biology but poorly understood as they are governed by non-equilibrium thermodynamics. Here, we bridge the gap between passive, close-to-equilibrium, and active, dissipative structures with chemically fueled phase separation. We observe that spherical, active droplets can transition into a new morphology—a liquid, spherical shell of droplet material. A spherical shell would be highly unstable at equilibrium. Only by continuously converting chemical energy, this dissipative structure can be sustained. We demonstrate the transition mechanism, which is related to the activation of a product outside of the droplet, and the deactivation within the droplets leading to gradients of droplet material. We characterize how far out of equilibrium the spherical shell state is and the chemical power necessary to sustain it. Our work suggests new avenues for assembling complex stable morphologies, which might already be exploited to form membraneless organelles by cells.

## Introduction

Spontaneous structure formation through self-assembly and phase separation (*1*) is essential in biology and engineering. These structures form spontaneously through the minimization of free energy leading to in- or close-to-equilibrium morphologies. Typical examples are structures formed by amphiphiles (*2*–*4*), nanoparticles (*5*), liquid crystals (*6*), peptides (*7*–*9*), and the demixing of immiscible liquids (*10*), which find application in healthcare (*11*), optoelectronics (*12*), and others (8). In contrast, dissipative structures can only form when a system is forcefully kept from reaching a free energy minimum by a continuous energy supply (*13*–*16*). Chemically fueled droplets are an example of a dissipative structure since their maintenance requires a continuous supply of free energy and matter (*17–23*). These structures are ubiquitous in biology but remain poorly understood, and synthetic models are rare (*24*).

In contrast, in- or close-to-equilibrium droplets (passive droplets) are well-understood (*25*–*30*). Such droplets form via phase separation in mixtures of immiscible liquids. They tend towards a spherical shape, corresponding to minimal interfacial surface energy. Minimizing this energy also drives coarsening through fusion and Ostwald ripening, leading to a single droplet (Fig. 1A).

**Figure 1.**
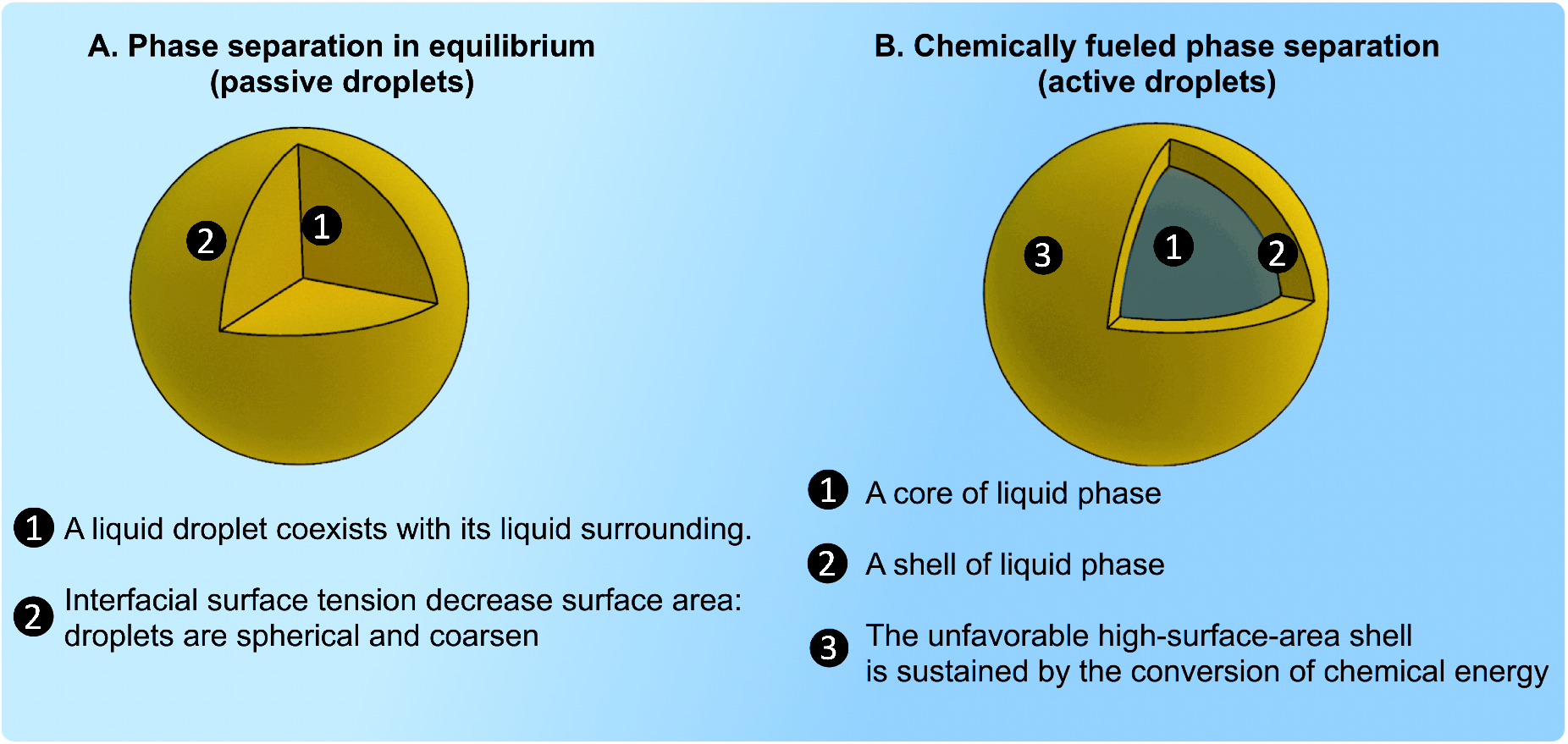
**A.** Phase separation close to thermodynamic equilibrium leads to spherical droplets. **B**. A new morphology is described in this work: a liquid, spherical shell.

In contrast to passive droplets, chemically fueled, active droplets are a new class of dissipative structures whose properties are governed by chemical reactions and diffusive fluxes. Theoretical studies on these structures show exciting behavior that results from their nonequilibrium nature, which includes inhibiting Ostwald ripening and thereby controlling their size control (*31*, *32*) and a spontaneous self-division (*33*, *34*). These behaviors require a non-equilibrium steady state which has so far been hard to archive under experimental conditions (*35*, *36*). Synthetic analogs are scarce to test these exciting behaviors, and methods to continuously fuel these droplets do not exist.

In this work, we describe how continuously fueled phase separation yields a new dissipative structure: a thin, spherical shell of phase-separated liquid (Fig. 1B). This is intriguing since the additional interface and, thus, the larger surface area compared to volume makes such spherical shells thermodynamically very unstable. The energy supply to sustain the thermodynamically unstable state comes from converting fuel to waste, leading to diffusive fluxes that keep the spherical shell stable. We show that this non-equilibrium spherical shell state forms due to an instability of the active droplet’s core and that the shell interfaces act as a pump for fuel-activated chemical components.

## Results and discussion

In our chemically fueled droplets, DIC (N,N’-diisopropylcarbodiimide) is the high ernegy molecule (fuel) that reacts with the C-terminal aspartic acid of a peptide (Ac-F(RG)3D-OH, precursor, Fig. 2A). It converts it into its corresponding cyclic anhydride (product, Fig. 2A). That anhydride has a half-life of 58 seconds (k_deact_.= 0.012 ± 0.009 s^-1^, Fig. S1, Table S5) before it is deactivated through hydrolysis. Upon activation, the peptide turns from zwitterionic to cationic with an overall charge of +3 (k_act_.= 0.17 ± 0.008 M^-1^s^-1^, Fig. S1, Table S5). In their short lifetime, the cationic, activated peptide can bind a polyanion (pSS, 400 monomer repeat units, 75kDa). The non-activated, zwitterionic precursor interacts weakly with the polyanion pSS (K_D, prec._ of 104.5 ± 11.4 μM, Fig S2). In contrast, the product interacts strongly with pSS (K_D, prod._ of 2.4 ± 1.2 μM, Fig. S2 and supplementary discussion 1). Thus, chemical fuel creates a population of short-lived activated building blocks that can interact with polyanions and phase separate into complex coacervate droplets.

**Figure 2.**
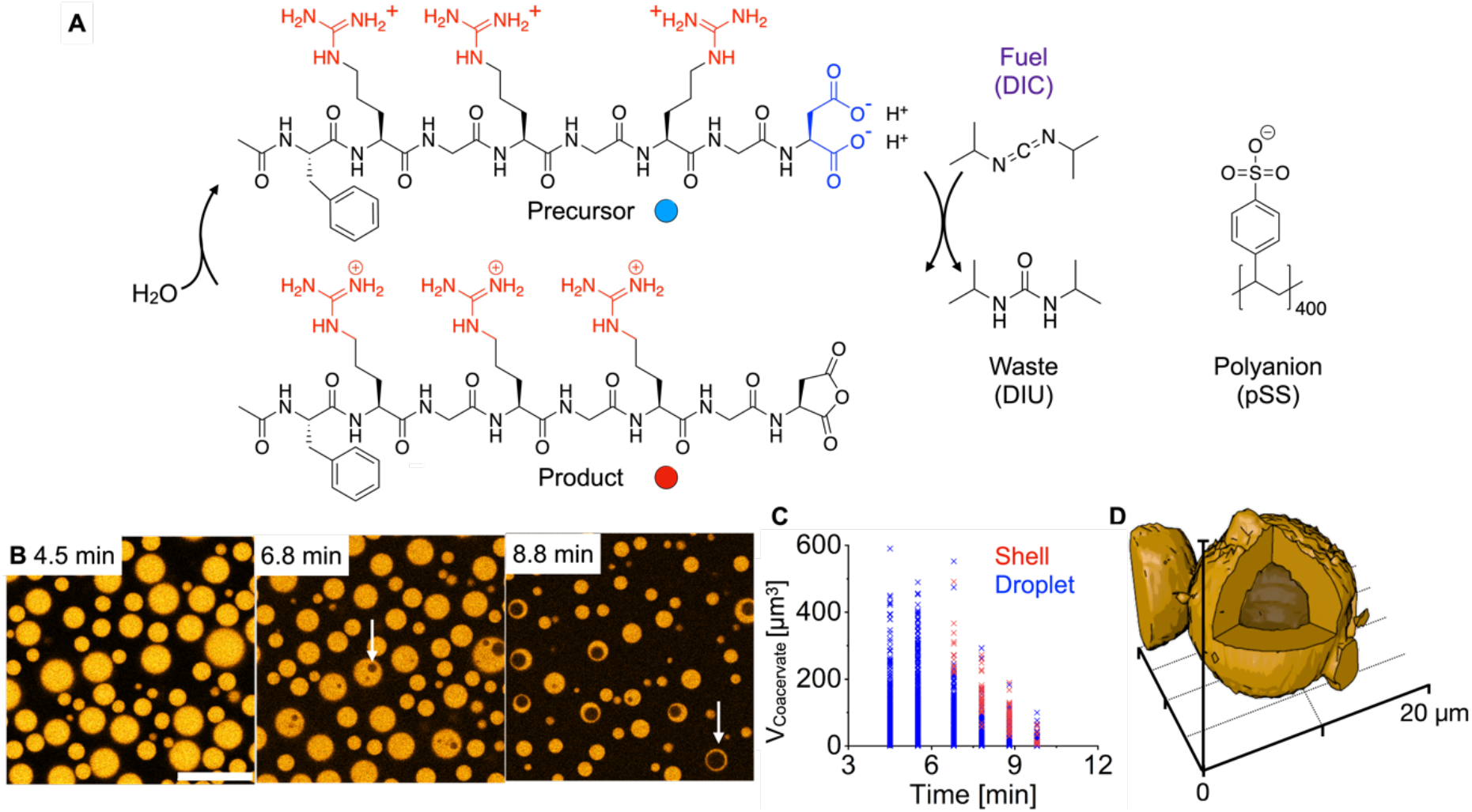
Chemically fueled droplets under batch-fueling. **A**. Chemical structures of the components involved in forming chemically fueled droplets and liquid shells. **B**. Confocal microscopy of a solution of 22 mM precursor, 12 mM pSS, and 0.1 μM sulforhodamine B in 200 mM MES buffer at pH 5.3 fueled with 20 mM DIC. After 7 minutes, the active droplets became unstable and swelled to form a spherical shell (white arrows). Scale bar 20 μm. **C**. The volume of each active droplet (blue) and active liquid shell (red) is shown for each time point. Larger active droplets formed spherical shells earlier than smaller active droplets. **D.** 3D reconstruction from confocal microscopy data shows the spherical nature of the shell. Conditions were 10 mM precursor, 5 mM pSS, and 0.1 μM sulforhodamine B in 200 mM MES buffer at pH 5.3 fueled with an excess of DIC (5μL) on top of 20μL of the sample. The diffusion of excess fuel into the sample leads to the formation of bigger droplets that are longer lived. The z-stack for the 3D reconstruction is imaged two hours after adding fuel.

When we fueled a solution of 22 mM peptide and 12 mM pSS (the concentration is expressed in repeat units) in 200 mM MES buffer with a batch of 20 mM fuel, we found that the sample turned turbid (Fig. S3A) due to the formation of micron-sized coacervate-based droplets as evidenced by confocal microscopy (Fig. 2B). These droplets fused, and, when the chemically fueled emulsion has depleted most of its fuel, the droplets decayed—a clear solution without droplets was obtained (Fig.S3A). In the dissolution process, after around 7 minutes, when the fuel ran low, we found that the droplets transitioned from a homogenous droplet to a spherical shell (Fig. 2B, Movie S1), a process referred to as vacuolization (*37*–*39*) which is also reminiscent of bubbly phase separation in active fluids (*40*). Moreover, we found that the larger the droplet, the sooner the droplet became unstable and formed a spherical shell (Fig. 2C and D).

If the solution was fueled with only 5 mM DIC, the droplets stayed smaller and did not form spherical shells during their dissolution (Fig. S3 B, Movie S2). Vacuole formation has previously been shown as a dissolution pathway of coacervate-based droplets (*18*, *41*, *42*). In our case, we hypothesize that these spherical shells form because when fuel runs low, the influx of new droplet material decreases, and the deactivation of the peptide in the core outcompetes the influx of new material. Thus, gradients of product material could form within the droplet, leading to the dissolution of the droplet’s core. A larger droplet would have a steeper gradient and thus dissolve sooner. This mechanism suggests that spherical shells could form and be sustained in steady state when fuel is provided continuously.

Continuously fueling soft matter to yield sustained non-equilibrium steady states is challenging because these solutions cannot be stirred without affecting the assemblies. Moreover, waste accumulation often poisons the self-assembling systems (*36*, *43*). Therefore, we developed new aqueous microreactors in which fuel can continuously diffuse (Fig. 3A and B). We made these aqueous microreactors by preparing a stable emulsion of water droplets in a fluorinated oil phase. The water droplets (microreactors) contained the peptide, buffer, and polyanion, *i.e*., all reagents except the fuel. The fluorinated oil contained the fuel, which diffused into the microreactors at the oil-water interface until a steady state of fuel and product was reached (Fig. 3A and S4A-B). Excitingly, the waste of the reaction cycle (DIU) partitioned preferentially outside the microreactors and crystallized in the fluorinated oil phase (Fig. S4A,C).

**Figure 3.**
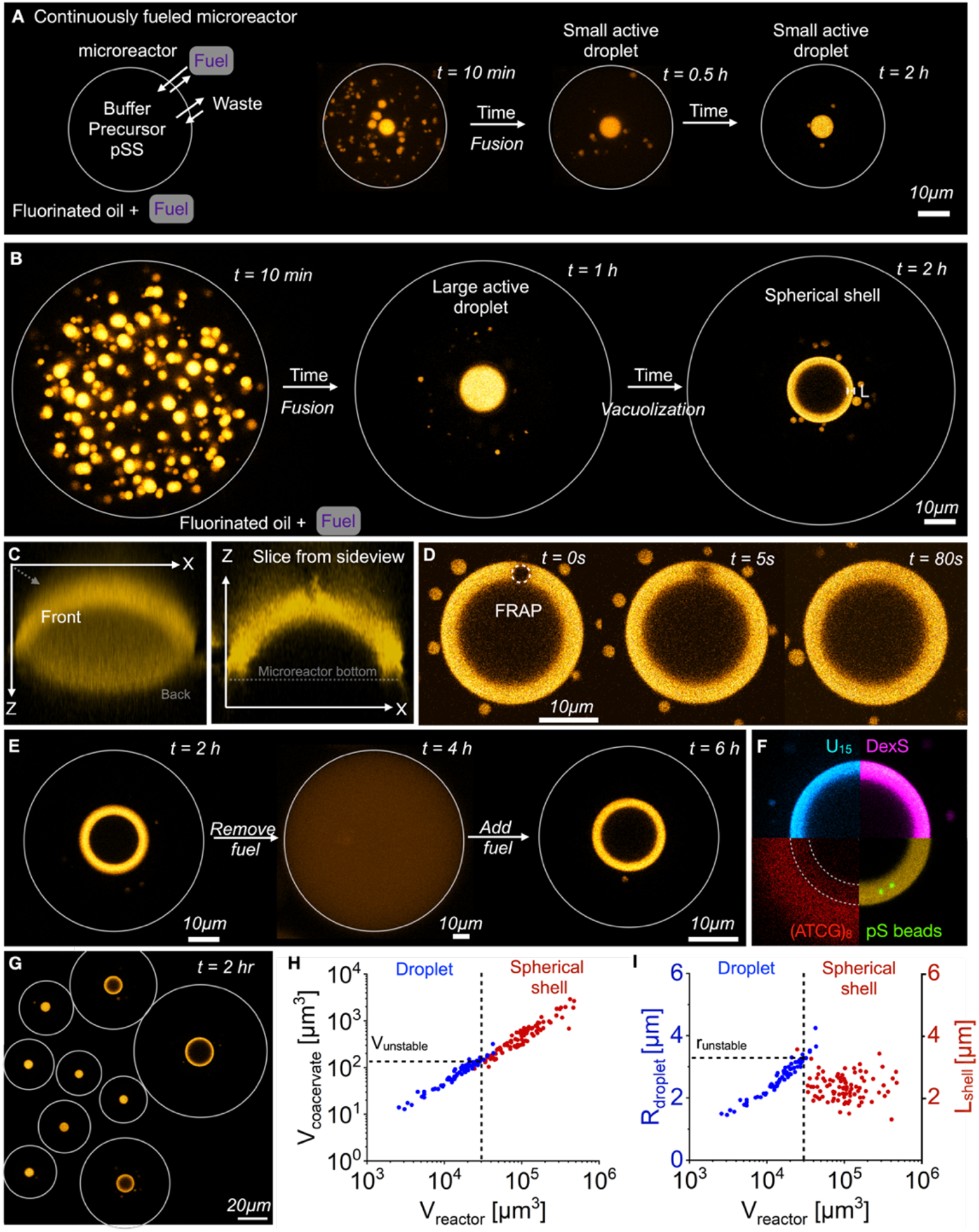
Spherical shells are a stable, non-equilibrium state. **A-B**. Experimental setup to form microreactors that continuously fuel active droplets. Surfactant-stabilized microfluidic droplets (microreactors, grey outline) containing 10 mM precursor, 5 mM pSS, and 0.1 μM sulforhodamine B in 200 mM MES buffer at pH 5.3 were embedded in a fluorinated oil which contained 0.5 M DIC to sustain the microreactor at a fuel concentration of 8.4 mM. Time-lapsed micrographs of a small (**A**) and large (**B**) microreactor in a steady state. The grey circle represents the size of the microreactor. The images for 10 min and 0.5 h in **A** and the image for 10 min in **B** are z-projections of the microreactor. All other images are from one Z-plane. **C.** A 3D projection of a spherical shell reconstructed from Z-stack imaging. The left image shows the projection of the average pixel intensity. The right image shows a slice in the XZ plane through the middle of the spherical shell. **D**. A FRAP study of the spherical shell demonstrates that the shell is liquid and dynamic. **E**. The microreactor was fueled with 0.5 M DIC in the oil phase. After the formation of spherical shells (2 h), the oil phase was replaced with oil containing no DIC. After a homogenous solution was obtained (4 h), the oil was replaced with 0.5 M DIC-containing oil. The grey circle represents the size of the microreactor. **F**. Partitioning of different fluorescently labeled molecules into the spherical shell. The dotted line represents the outlines of the spherical shell. **G**. A macroscopic view of multiple microreactors shows that large reactors formed spherical shells while small microreactors contained droplets. The center Z-plane of each of the individual droplets and shells is projected. The grey circle represents the size of the microfluidic reactor. **H.** Above a critical reactor volume, active droplets transformed into spherical shells. **I**. Spherical shells had a similar thickness L_shell_, which was within the range of r_unstable_.

For the experimental conditions of Fig. 2 (0.5 M DIC in the oil phase), a concentration of 8.4 mM DIC in the aqueous phase was measured (Fig. S4D). Moreover, the concentration of fuel in the fluorinated oil controlled the fuel concentration inside the microreactor, too (Fig. S4D). Thus, this new continuous fueling method allows us to continuously fuel microreactors at various steady states and simultaneously avoid the challenge of accumulating waste. Z-stacks of the microreactors can be acquired with a confocal microscope. When projected onto one plane, each coacervate droplet can be counted, and its volume measured (Fig. 3A). Finally, with our method, a large number of microreactors of volumes ranging from thousands to millions of μm^3^ are made in one experiment (Fig. 3G).

Within seconds after preparing the microreactors, coacervate droplets nucleated and grew through fusion (Fig. 3A). After about 10 minutes, the total volume of all combined droplets within one microreactor reached its maximum. It stayed constant over the entire time of the experiment, a strong indication that a steady state in activation and deactivation was reached (Fig. S4E-F, Movie S3). Within 30 minutes, all droplets fused until one large active droplet remained. We identified two possible behaviors of the active droplet: in small microreactors, the active droplet sunk to the bottom of the reactor, slightly wetted the reactor floor and remained stable for the entire observation time (Fig. 3A, Fig. S5A, and S6A-B).

In large microreactors, the active droplet was also larger. These droplets sunk and slightly wetted the reactor floor (Fig. 3B). However, after about 2 h, its core became unstable. Remarkably, the droplet transitioned and left behind a shell of droplet material homogenous in thickness (Fig. 3C) that was stable for several hours (Fig. S6A-C). As the original droplet had wetted the floor, we did not obtain a spherical shell as for the experiments in batch-fueling. Instead, we observed a half dome of phase-separated material (Fig. 3C). If we performed the same experiments with passive droplets, we did not observe the formation of any spherical shells (Fig. S7, Fig. S5B-C). We measured the product’s diffusion coefficient within the active droplets and the active spherical shells, D_prod._, by fluorescence recovery after photobleaching (FRAP) to be 0.03 μm^2^/sec (Fig. 3E and S8B,D, Table S4). FRAP confirmed the shell was liquid which was further corroborated when we exchanged the fluorinated oil with fluorinated oil without fuel. The spherical shell rapidly dissolved until all droplet material had decayed (Fig. 3E). When the oil was exchanged with DIC-loaded oil, droplets reappeared that fused. Still, due to extensive wetting of the newly formed active droplets, only a small fraction of the microreactors yielded into spherical shells.

Next, we tested the permeability and partitioning of the shell by adding fluorescently labeled anionic molecules and nanoparticles: a U15 RNA oligomer, dextran sulfate, carboxy-terminated polystyrene particles, and a DNA oligomer (ATCG)_8_ (Fig. 3F). All partitioned well into the shell except for (ATCG)_8_ which showed a lower fluorescence in the shell than in the dilute phase. Additionally, both U_15_ and dextran sulfate showed similar fluorescence within the interior of the spherical shell compared to the dilute phase outside (Fig. S9A-B), whereas (ATCG)_8_ was excluded from the interior of the spherical shells (Fig. S9C). This suggests that the shell acts as a permeable membrane to molecules that partition well, but not ones that hardly partition or large particles that do not leave the spherical shell (*44*).

We quantified the relationship between the reactor volume and droplet behavior. We first confirmed the volume of the droplet material scales linearly with the reactor volume (Fig. 3G and H). Simply put, a larger microreactor contains more precursor molecules and produces more product molecules. This dataset revealed that a shell could be expected when the microreactor was larger than roughly 30.000 μm^3^ (Fig. 3H). This microreactor volume corresponds to a coacervate-based droplet volume of 150 μm^3^. In other words, shells were formed when the microreactor was sufficiently large to form coacervate-based droplets larger than 150 μm^3^ corresponding to the threshold radius r_unstable_ = 3.3 μm (Fig. 3I). Additionally, we found that the thickness of the spherical shells L_shell_ was quite constant (L_shell,exp_. = 2.4 ± 0.4 μm) over a large range of total phase-separated material (V_coacervate, shell_ = 150 to 3000 μm^3^).

To understand the mechanism of the spherical shell formation, we measured the product’s and fuel’s partition coefficients in the coacervate-based droplets with a spin-down assay. We found that the fuel only weakly partitions in the droplets (K_fuel_ = 1.4 ± 1.7, Table S3, supplementary discussion 2). Thus, most of the fuel remains outside the droplets. In contrast, the product partitions strongly inside the droplets (K_product_ = 3360 ± 1645, Table S2). We conclude that activation predominantly occurs outside, whereas deactivation occurs inside the droplets. The spatial separation of these reactions leads to a diffusion gradient of product that builds up from the outside of the droplet towards its core (Fig. 4A). In other words, the greater the radius of the active droplet, the more its core is depleted from the product. It thus destabilizes, leading to the core’s instability and the transition into a spherical shell (Fig. 4B and Fig. S10, Movie S4).

**Figure 4.**
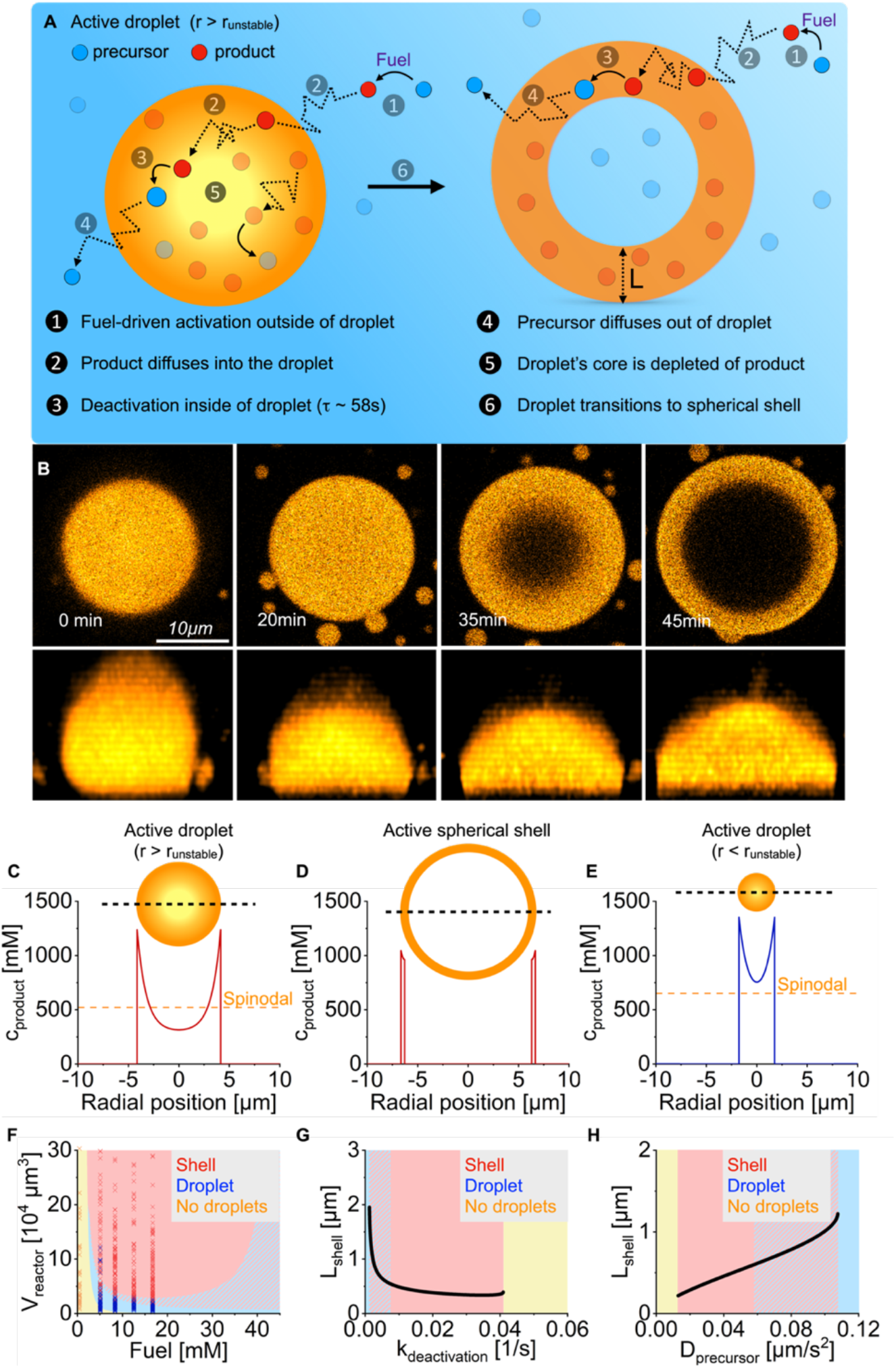
The mechanism of spherical shell formation. **A.** Schematic representation of the mechanism of spherical shell formation. **B**. Confocal micrograph timelapse series of an active droplet with a critical radius larger than r_unstable_. Over 20 minutes, the droplet wetted the microreactor’s bottom and transitioned into a spherical shell. The scale bar represents 10 μm. The upper row of images shows the XY-plane close to the bottom of the reactor. The bottom row of images shows the XZ-projection. **C-E**. Concentration profiles of a large chemically fueled droplet with r > r_unstable_ (**C**) before transitioning into a spherical shell, a spherical shell (**D**), and a small chemically fueled droplet with r < r_unstable_ (**E**). The insets show a scheme of the droplet and spherical shell. **F**. The system’s behavior as a function of steady-state concentration fuel and reactor volume. The shaded areas represent the stable state calculated by the model. The red-blue shaded area represents the coexistence of stable droplets and shells. The markers show the phase-separated state of the experimental data. **G-H**. The shell thickness L_shell_ for a microreactor with a radius of 25 μm as a function of varying deactivation rate constants (**G**) or precursor diffusion constants (**H)**.

To quantitatively verify that hypothesis, we developed a non-equilibrium thermodynamics-based model that describes a droplet in the center of a spherical reactor with the fuel and waste concentrations maintained at the reactor’s boundary. We used experimentally determined steady-state concentrations, rate constants, diffusion, and partitioning coefficients (Table S1-5). To provide a theory with the minimal physiochemical ingredients, we do not describe the poly-anion and the wetting of the spherical shell on the microreactor wall. We experimentally determined the concentrations of the three components in the phase diagram (see supplementary discussion 2), which was then fitted by a mean-field Flory Huggins model showing good agreement (Fig. S12).

We used our model to calculate the product’s concentration profile from a droplet core to its boundary (Fig. 4C). We used a droplet with a 5 μm radius which is greater than r_unstable_ (Fig. 3I). The concentration decreases from 1200 mM at the droplet interface to 450 mM at the core, which is depleted of droplet material because of the deactivation. The core’s concentration is far below the spinodal decomposition concentration. Thus, the droplet core cannot be stable, *i.e*., a new domain more similar to the outside nucleates at the core and grows, leaving behind a spherical shell of droplet material (Fig. 4D). A droplet with a radius smaller than r_unstable_ has a core in which the product concentration remains above the spinodal decomposition concentration and is thus stable (Fig. 4E).

Strikingly, we found that the model predicted r_unstable_ to be 2 μm, *i.e*., close to the experimentally determined 3.3 μm. Further calculations created a diagram that predicts a droplet’s behavior based on its microreactor size and fuel concentration (Fig. 4F). At low steady-state fuel concentrations, insufficient droplet material is activated to form a stable droplet. Above 5 mM of fuel, phase separation is observed and a stable droplet or a spherical shell is found depending on the microreactor size.

Surprisingly, beyond a steady-state fuel concentration of 20 mM, large reactors produced stable droplets again. As the fuel concentration increases, its concentration inside the droplets also increases. Peptide re-activation inside the droplet weakens the concentration gradient of the product inside (Fig. S15). According to our model, the non-equilibrium steady states of an active spherical shell and an active droplet can coexist (blue-red shaded area). Some of these theoretical calculations could be experimentally verified. We changed the fluorinated oil’s fuel concentration to tune the microreactors’ fuel concentration between 0.4 and 16.8 mM (Fig. 4F, Fig. S11, Fig. S4D) to find agreement with the predicted microreactor sizes for the formation of spherical shells, droplets, and the homogenous phase.

In line with the experiments, the model also predicted that the thickness of the shell (L) is relatively constant with varying reactor sizes, despite it decreasing slightly towards bigger reactor volumes (Fig. S16). However, the absolute value is lower than the measured one (L_theory_ ~ 0.3 - 0.6 μm). Using the model, we calculated that L initially decreases rapidly with an increasing deactivation rate constant (k_deact._) but is then quite stable over a broad range of k_deact._. However, if the product is too short-lived, no droplets were found (Fig. 4G). In contrast, increasing the product’s diffusion constant made the shell thicker*, i.e*., the activated product can diffuse further in its short lifetime leading to a weaker gradient. Also, here, a too-fast diffusing product led to no droplets (Fig. 4H).

Finally, we calculated the free energy and the chemical work needed to produce spherical shells (see method section R). The free energy difference between a spherical shell and a droplet of identical volume is reflected in the additional interface. For an assumed surface tension (*γ*) of 75 μN/m, the additional free energy cost (F_surface_) of a spherical shell with an inner radius (R_in_) of 2 μm equals 4 fJ. Based on the equilibrium constant of the acid-anhydride equilibrium (*45*), we could calculate a free energy difference between the precursor and the product of about 10 k_B_T (F_act_ = 80 nJ). Therefore, the additional surface energy is negligible compared to the free energy needed for activating the precursor to the product to sustain the spherical shell. From the model, we calculated the number of chemical cycles to sustain the steady state to be 6·10^4^/(μm^3^ s). Combined with the free energy difference of the cycle, the total free energy turned over per time, and volume is J_tot_ = 0.275 W/L. Simply put, this power is supplied to maintain the product population in the activated state.

Nevertheless, what fraction of this power is used to maintain the spherical shell morphology? The spherical shell results from activation dominating deactivation outside, while inside, there is net deactivation. This imbalance of chemical rates gives rise to a net diffusive influx of product and efflux of the precursor at the shell interfaces, J_int_. The flux of the product multiplied by the activation energy of 10 k_B_T corresponds to the power to keep the shell interface steady and prevent the relaxation to the active droplet or even the homogenous state. Thus, the imbalance of chemical rates between the two coexisting phases gives rise to an energy pump. We find that the power of this pump is about J_int_ = 0.198 W/L (see method section R). Compared to the supplied chemical power, this yields an efficiency, J_int_/J_tot,_ of about 72%. The efficiency is lower than 100% because the deactivation outside as well as the re-activation inside the shell do not contribute to the flux though the interface.

## Conclusions

In our work, we described a new microreactor to sustain chemically fueled droplets in a steady state. Here, a precursor is activated to form droplets at the expense of chemical fuel. We found a transition from an active droplet to a liquid spherical shell state which is the result of a spinodal instability of the active droplet’s core. Due to this instability, the core material gets spontaneously pushed out of the droplet’s core leading to the formation of an additional inner interface. The resulting spherical shell is a new non-equilibrium steady state. We showed that the spherical shell acts as an energy pump transporting activated product molecules into the shell and deactivated precursor molecules toward its outside. Quantifying the free energy turnover, we found that this pump is rather efficient, *i.e*., most fuel-activated components are used to stabilize the additional spherical shell interface.

The power consumption of a fuel-driven spherical shell of 0.275 W/L is smaller but comparable to the power consumption of living cells, which is roughly 1 W/L. One reason is that the concentration of fuel (ATP in cells) and the energy it liberates upon hydrolysis are in the same range as our synthetic system (*46*). Interestingly, reminiscent morphologies to spherical shells were also observed in various cell types (*37*, *47*).

## Acknowledgments

The BoekhovenLab is grateful for support by the TUM Innovation Network - RISE funded through the Excellence Strategy and the European Research Council (ERC starting grant 852187). This research was conducted within the Max Planck School Matter to Life supported by the German Federal Ministry of Education and Research (BMBF) in collaboration with the Max Planck Society. Funded by the Deutsche Forschungsgemeinschaft (DFG, German Research Foundation) under Germany’s Excellence Strategy - EXC-2094 – 390783311. C. Weber acknowledges the European Research Council (ERC) under the European Union’s Horizon 2020 research and innovation programme (Fuelled Life, Grant Number 949021) for financial support.

## Supplementary Materials

Materials and Methods

Supplementary Text

Figs. S1-14

Tables S1-7

Movies S1-4

## Notes

### Competing Interest Statement

The authors have declared no competing interest.

