## Supplementary Information for "Liquid spherical shells are a non-equilibrium steady state"

*equal contribution

[+]corresponding author

### Table of Contents


I. Material and Methods

- (A) Materials
- (B) Standard sample preparation
- (C) Nuclear magnetic resonance spectroscopy (NMR)
- (D) High-performance liquid chromatography (HPLC)
- (E) Fluorescence spectroscopy
- (F) Determination of rate constants of the chemical reaction cycle
- (G) Kinetic model
- (H) Isothermal titration calorimetry (ITC)
- (I) Determination of the partitioning and concentration of molecules inside and outside of the droplet phase
- (J) Determination of total droplet volumes
- (K) Confocal fluorescence microscopy and droplet analysis
- (L) Fluorescence recovery after photobleaching
- (M) Method to continuously fuel active droplets
- (N) Estimating the concentration of product and fuel in the microfluidic droplet
- (O) Thermodynamic model for the experimental phase diagram
- (P) Sharp interface model for the kinetics of active droplets and active spherical shells
- (Q) Parameter values used in numerical calculations
- (R) Calculations of free energies and free energy rates

II. Supporting Discussion

III. Supporting Tables

IV. Supporting Figures

V. Supporting Movies

VI. References

### I. Material and Methods

**(A) Materials.** All solvents were purchased in analytical grade from Sigma Aldrich and used without further purification. *N,N*-dimethyl formamide (DMF), Fmoc-R(Pbf)-OH, Fmoc-D(OtBu)-OH, Fmoc-G-OH, Fmoc–N(Trt)-OH), *N,N'*-diisopropylcarbodiimide (DIC), ethyl cyano(hydroxyimino)acetate (Oxyma, Nova-biochem©), Wang resins (100- 200 mesh, 0.4-0.8 mmol/g), Rink Amide resins (100-200 mesh, 0.4-0.8 mmol/g), 4-(dimethylamino)-pyridine (DMAP), trifluoroacetic acid (TFA, 99%), piperidine (99%), triisopropylsilane (TIPS), *N,N'*-diisopropylethylamine (DIPEA), 4-chloro-7-nitrobenzofurazan (NBD-Cl, 98%), polystyrene sulfonate (pSS, 75kDa, 18wt% in water), sulforhodamine B, 4-morpholineethanesulfonic acid (MES) buffer, carboxylate-modified polystyrene beads (fluorescent yellow-green, 1 μm mean particle size), Cy3-labeled $(ATCG)_8$ were all purchased from Sigma-Aldrich and used without any further purification unless indicated otherwise. Peptides Ac-F$(RG)_3$D-OH and Ac-F$(RG)_3$N-$NH_2$ were purchased from CASLO Aps. Peptides NBD-G$(RG)_3$N-$NH_2$ and NBD-G$(RG)_3$D-OH were synthesized based on a published procedure (*1*). Cy5-pSS (710 kDa) was synthesized using a published procedure (*2*). Cy3-$U_{15}$ was purchased from biomers.net GmbH. Cy3-dextran sulfate (40 kDa) was purchased from CD Bioparticles. 2w% 008-Fluorosurfactant in Novec 7500 was purchased from RAN Biotechnologies. Novec 7500 was purchased by Iolitec.

**(B) Standard sample preparation.** Stock solutions of Ac-F$(RG)_3$-D-OH (precursor, 300 mM), pSS (41 mM, according to monomer units), and MES (650 mM) were prepared by dissolving the respective amount of each component in MQ water and adjusting the pH to 5.3. Stock solutions for Ac-F$(RG)_3$N-$NH_2$ (product*), and the fluorescent dyes were also prepared in MQ water but without pH adjustment. All stocks were filtrated with a syringe filter (PTFE, 0.2μm pore size). For active droplets fueled with a batch of fuel: 5-20 mM DIC (fuel) was added to a solution (20-500 μL) of precursor and pSS in 200 mM MES at pH 5.3 and the solution was mixed with a pipette. For active droplets fueled with an excess of fuel: 5 μL of DIC is added to a solution (20 μL) of 10 mM precursor and 5 mM pSS in 200 mM MES at pH 5.3. The sample is mixed with a pipette and residual DIC above the solubility limit is left as a reservoir on top of the aqueous solution. For active droplets fueled continuously, a solution (5 μL) containing precursor and pSS in 200 mM MES at pH 5.3 was mixed with perfluorinated oil (Novec 7500, 50 μL) containing the fluorosurfactant and 0.025-1.5 M DIC. For passive droplets that are not fueled, pSS was added to a solution containing precursor and product* in 200 mM MES at pH 5.3 and the solution was mixed with a pipette.

**(C) Nuclear magnetic resonance spectroscopy (NMR)**. $^{1}$H-NMR spectra were recorded with 16 or 64 scans at room temperature on Bruker AVHD 300, AVHD400, and AVHD500 spectrometers. Chemical shifts are reported in parts per million (ppm) relative to the signal of the deuterated solvent $D_2O$ ($\partial$ = 4.7 ppm). All measurements were performed in triplicate (N=3) at 25 °C.

**(D) High-performance liquid chromatography (HPLC)**. High-pressure liquid chromatography was performed using analytical HPLC (ThermoFisher, Vanquish Duo UHPLC, and Thermofisher Dionex Ultimate 3000) with a Hypersil Gold C18 column (100 x 3mm, 250 x 4.6mm). Separation was performed using a linear gradient of acetonitrile (2 to 98%) and water with 0.1% TFA and the chromatogram was analyzed using detectors at 220 nm and 254 nm. The data was collected and analyzed with the Chromeleon 7 Chromatography Data System Software (Version 7.2 SR4). All measurements were performed in triplicate (N=3) at 25 °C.

**(E) Fluorescence spectroscopy.** Fluorescence spectroscopy was performed on a Jasco spectrofluorimeter (Jasco FP-8300, SpectraManager software 2.13) with external temperature control (Jasco MCB-100). All experiments were performed in triplicate (N=3) at 25 °C.

**(F) Determination of rate constants of the chemical reaction cycle.** The concentration profiles of the precursor and its corresponding anhydride (product) were determined by HPLC, whereas NMR quantified fuel and waste.

Quantification of precursor and product: due to the instability of the product, a quenching technique was used with an amine that converts the product into a quantifiable amide (*1, 3, 4*). To a solution (145 µL) of 10 mM precursor in 200 mM MES at pH 5.3 was added a solution (5 µL) of 5-20 mM DIC in acetonitrile in an HPLC vial. After each time point, 10 µL of the reaction mixture was added to 20 µL of an aqueous solution of ethylamine (400 mM). The resulting clear solution was then injected into the HPLC, and the concentration of the precursor and product was calculated from the resulting peak integrals.

Quantification of fuel and waste: 5-20 mM DIC was added to a solution (1 mL) of 200 mM MES at pH 5.3 and vortexed for 30 s to dissolve all DIC. The sample also contained 10 vol% $D_2O$ and 80 mM acetonitrile (ACN) as a reference. 10 mM precursor was added to 500 µl of this solution. The solution was vortexed and added to an NMR tube. $^{1}$H-NMR measurements were performed on a 300 MHz NMR every 5 min until no more DIC was detected. The concentrations of fuel and waste were then calculated from the peak integrals, which were compared with the acetonitrile reference. The chemical shifts of the compared signals were

[1]H NMR (300 MHz, $D_2O$): ACN $\delta$ (ppm) = 2.07 (s, 3H, CH3), DIC $\delta$ (ppm) = 1.22-1.23 (d, 12H, CH3), DIU $\delta$ (ppm) = 1.09-1.10 (d, 12H, CH3).

The obtained data from the HPLC and NMR experiments were then fitted via a kinetic model in python.

**(G) Kinetic model.**

The reaction cycle in a homogeneous system is modeled according to the following mechanism:

$$F \overset{k_0}{\rightarrow} W$$

$$Ac + F \overset{k_1}{\rightarrow} I$$

$$I \overset{k_{An}}{\longrightarrow} An + W$$

$$I \overset{k_{Ac}}{\longrightarrow} Ac + W$$

$$An \overset{k_2}{\rightarrow} Ac$$

Where Ac is the dicarboxylic acid, F is the fuel, I is the intermediate O-acylurea, W is the waste and An is the anhydride.

The mechanism translates into the following set of differential equations:

$$\begin{cases} \frac{d[Ac]}{dt} = -k_1 \cdot [Ac] \cdot [F] + k_2 \cdot [An] + k_{Ac} \cdot [I] \\ \frac{d[F]}{dt} = -k_1 \cdot [Ac] \cdot [F] - k_0 \cdot [F] \\ \frac{d[W]}{dT} = +k_0 \cdot [F] + k_{Ac} \cdot [I] + k_{An} \cdot [I] \\ \frac{d[An]}{dt} = +k_{An} \cdot [I] - k_2 \cdot [An] \\ \frac{d[I]}{dt} = +k_1 \cdot [Ac] \cdot [F] - k_{An} \cdot [I] - k_{Ac} \cdot [I] \end{cases} \tag{1}$$

We then applied steady-state approximation to obtain:

$$\frac{d[I]}{dt} = +k_1 \cdot [Ac] \cdot [F] - k_{An} \cdot [I] - k_{Ac} \cdot [I] \approx 0 \Rightarrow [I] \approx \frac{k_1 \cdot [Ac] \cdot [F]}{k_{An} \cdot (\frac{k_{Ac}}{k_{An}} + 1)}$$

We called $\frac{k_{Ac}}{k_{An}} = K$ and used the relation above in the set of differential equations to obtain:

$$\begin{cases} \frac{d[Ac]}{dt} = -k_1 \cdot [Ac] \cdot [F] + k_2 \cdot [An] + \frac{K \cdot k_1 \cdot [Ac] \cdot [F]}{(K+1)} \\ \frac{d[F]}{dt} = -k_1 \cdot [Ac] \cdot [F] - k_0 \cdot [F] \\ \frac{d[W]}{dT} = +k_0 \cdot [F] + k_1 \cdot [Ac] \cdot [F] \\ \frac{d[An]}{dt} = +\frac{k_1 \cdot [Ac] \cdot [F]}{(K+1)} - k_2 \cdot [An] \end{cases} \tag{2}$$

Experimental data were fit to equation system (2) using a custom program in Python 3 (kinmodel, https://github.com/scotthartley/kinmodel) previously published by the group of Hartley and applied to similar systems (*5*). To calculate concentrations of precursor and product under continuous fueling, the change in fuel concentration was set to 0 ($\frac{d[F]}{dt}$=0).

**(H) Isothermal titration calorimetry (ITC).** ITC experiments were performed with a MicroCal PEAQ-ITC from Malvern Pananalytical. All experiments were performed at 25°C. The following conditions were used: pSS 75kDa (1 mM sulfonate units in MES 200 mM, pH 5.3) was titrated with the precursor (15 mM in 200 mM MES, pH 5.3): 25 injections, 1.5 µL/inj. pSS 75kDa (0.05 mM sulfonate unit in 200 mM MES, pH 5.3) was titrated with the product* (1.5 mM as charged units, 0.5 mM as molecular concentrations in 200 mM MES, pH 5.3): 25 injections, 1.5 µL/inj. All experiments were performed in triplicate (N=3). For each experiment, a control was performed by titrating the corresponding amount of peptide (precursor or product*) in 200 mM MES buffer (pH 5.3) in the absence of pSS. Data were fitted using a non-linear least squares algorithm provided with the PEAQ-ITC Analysis software.

**(I) Determination of the partitioning and concentration of molecules inside and outside of the droplet phase.** To determine the amount of precursor, product*, and fuel in the dilute phase and inside passive droplets for the phase diagram, we measured their concentrations using HPLC, fluorescence spectroscopy, and NMR.

Precursor: we quantified the fraction of the precursor that remained in the dilute phase of passive droplets by means of HPLC. 5 mM pSS was added to a solution containing 10-x mM precursor and x mM product* (total peptide concentration = 10 mM) or 20 mM precursor (without product*) in 200 mM MES at pH 5.3. The turbid suspension (V = 150 µL) was vortexed, incubated for 5 min, and then centrifuged for another 15 min at 20,412×g. 100 µL of the supernatant was removed and added into an HPLC inlet. 1 µL of an aqueous solution of NaCl (4 M) was added to dissolve residual turbidity. The resulting clear solution was injected into the HPLC and compared to a solution containing 10-x mM precursor or for 20 mM precursor without pSS to calculate the fraction of precursor that remained in the supernatant.

Product*: we quantified the fraction of product* as a measure for the anhydride product that remained in the dilute phase of passive droplets by means of fluorescence spectroscopy because its concentration in the supernatant could not be determined accurately via HPLC (peak overlap with precursor, low fraction remaining in supernatant). Therefore, we used NBD-G(RG)$_3$N-NH$_2$ (fluorescent analog of product*) which partitions similarly into the droplet phase as product*.(*2*) To a solution of 10-x mM precursor and x mM product* (total peptide concentration = 10 mM) or 2 mM product* (without precursor) in 200 mM MES at pH 5.3 with 1 µM NBD-G(RG)$_3$N-NH$_2$ were added 5 mM pSS. The turbid suspension (V = 150 µL) was vortexed, incubated for 5 min, and then centrifuged for another 15 min at 20,412×g. 100 µL of the supernatant was removed and added into an Eppendorf tube. 1 µL of an aqueous solution of NaCl (4 M) was added to dissolve residual turbidity. The fluorescence of the sample was then measured on the fluorimeter (Excitation at 467 nm, emission at 526 nm). To account for the dependence of the fluorescence of dyes on their environment (*6*), we prepared an identical sample without NBD-G(RG)$_3$N-NH$_2$ and added the supernatant (96 µL) to an Eppendorf tube containing 1 µL of an aqueous solution of NaCl (4 M). To the clear solution, we then added 1 µM NBD-G(RG)$_3$N-NH$_2$ (4 µL) as in the previous sample and measured its fluorescence intensity. The intensity ratio between these 2 samples was therefore the fraction of the fluorescent molecules that remained in the supernatant.

Fuel: we quantified the fraction of fuel that partitioned into the droplet phase of passive droplets by means of NMR. To avoid the reaction of precursor and fuel, passive droplets consisting only of product* and pSS were used. 10 mM DIC (1.55 µl) was added to a solution (1 mL) containing 2 mM product* and 5 mM pSS in 200 mM MES at pH 5.3. The solution was vortexed and incubated for 5 min. The sample was centrifuged for 5 min at 20,412×g and the supernatant was carefully removed. The residual coacervate phase (about 1 µL) was dissolved in a solution (50 µL) containing 400 mM NaCl, 560 mM borate buffer at pH 10, 80 mM ACN as standard, and 20 vol% $D_2O$. The high pH prevents further hydrolysis of DIC between sample preparation and measurement. the DIC concentration was determined by $^1$H-NMR spectroscopy. Blank measurements without droplets were performed to account for residual DIC containing dilute phase on the walls of the sample containers after the removal of the supernatant. As a blank, 10 mM DIC was added to a solution (1 mL) containing 200 mM MES at pH 5.3. The blank solution without droplets was treated similarly to the solution with droplets. The sample was centrifuged for 5 min at 20,412×g and the supernatant was carefully removed. A solution (50 µL) containing 400 mM NaCl, 560 mM borate buffer at pH 10, 80 mM ACN as standard, and 20 vol% $D_2O$ was added and the DIC concentration was determined by $^1$H-NMR spectroscopy. The concentrations of fuel were then calculated from the peak integrals which were compared with the acetonitrile reference and corrected by the blank. The chemical shifts of the compared signals were $^1$H NMR (500 MHz, $D_2O$): ACN $\delta$ (ppm) = 2.07 (s, 3H, CH3), DIC $\delta$ (ppm) = 1.22-1.23 (d, 12H, CH3).

Combined with the total volumes of the droplet pellets (see method below), we calculated the partitioning of the individual molecules in passive droplets (Table S1-3). The resulting error bars of the concentration inside the droplets ($c_{in}$) and the partitioning coefficient (K) represent the accumulated standard deviation from the experiments measuring the fraction of molecules in the supernatant ($c_{out}$) and the total volumes of the coacervate pellets.

**(J) Determination of total droplet volumes.** To calculate the partitioning of the precursor, product*, and fuel in the droplet phase for the phase diagram, the total droplet volume was estimated via a centrifugation assay. For that, samples (200 µL) of passive droplets containing 10-x mM precursor and x mM product* (total peptide concentration = 10 mM), 20 mM precursor (without product), or 2 mM product* (without precursor) in 200 mM MES at pH 5.3 with 15 µM sulforhodamine B (for visualization) were prepared. After the addition of 5 mM pSS, the turbid suspensions were incubated for 5 min and then centrifuged for another 15 min at 20,412×g. The volume of the droplet pellets (0.4-2 µL) was then compared to size standards visually (Table S1-3).

**(K) Confocal fluorescence microscopy and droplet analysis.** A lightning SP8 confocal microscope (Leica) with a 63x water immersion objective (1.2 NA) was used to analyze the coacervates in bulk and in the microfluidic reactors (microreactors). Sulforhodamine B was added to track the coacervates via fluorescence, and the dye was excited at 552 nm and imaged at 565-700 nm with a HyD detector. The pinhole was set to 1 Airy unit.
Coacervate-based droplets and coacervate-based spherical shells were analyzed with ImageJ (Fiji). For the analysis, images of coacervate-based droplets were thresholded with the moments thresholding algorithm and a gaussian blur was applied. Subsequently, the images were analyzed with the analyze particle plugin in ImageJ. The used parameters were size = 5 $\mu m^2$-infinity, circularity= 0-1, and all images were analyzed once with holes included and once without holes. The thickness of the coacervate-based spherical shells was determined by subtracting the vacuole's radius from the droplet's radius with holes. The volume of coacervate-based spherical shells was calculated assuming a spherical shape and considering the half-sphere shape due to wetting at the interface of the microreactor and the oil. The volume of coacervate-based droplets without vacuoles was calculated assuming a spherical droplet.

**(L) Fluorescence recovery after photobleaching.** The diffusivity of the molecules inside active droplets was measured via spot bleaching. Measurements were performed in microfluidic reactors. Coacervate-based droplets were stained with NBD-G$(RG)_3$D-OH (fluorescent analog of precursor) or NBD-G$(RG)_3$N-$NH_2$ (fluorescent analog of product*) which was excited at 488 nm and imaged at 565-635 nm with a PMT detector. A spot size of r = 1 µm in radius was bleached in all experiments. Recovery data were normalized through double normalization with the following equation (7):

$$F(t) = \frac{(T_0 - B_o)(I_t - B_t)}{(T_t - B_t)(I_0 - B_0)}$$

Here F(t) represents the normalized fluorescence intensity which is calculated from the average intensity of 3 ROIs. It represents the average intensity of the bleached ROI, $T_t$ represents the average intensity of an unbleached ROI within the bleached coacervate and $B_t$ represents the average intensity of a ROI without any coacervates. The fluorescence recovery F(t) is then given by the equation (*8*):

$$F(t) = F_{\infty}\exp\left[-\frac{2}{1+\left(\frac{8tD}{a^2}\right)}\right]$$

with $F_{\infty}$ representing the fluorescence at full recovery, *a* representing the radius of the bleached area, and *D* representing the translational diffusion coefficient of the fluorescent probe.

**(M) Method to continuously fuel active droplets.** Surfactant-stabilized water in oil droplets was produced using 1 % 008-FluoroSurfactant in 3M HFE7500 as the oil phase. To form microreactors of varying size, 5 µL of a solution containing 10 mM precursor and 5 mM pSS in 200 mM MES at pH 5.3 were added to 50 µL of the oil phase in a 200 µL Eppendorf tube. Active droplets: For the preparation of active droplets, fuel was added to the oil phase before the addition of the sample containing the precursor and pSS. Snipping of the centrifugal tube resulted in the formation of microreactors with a random size distribution. The microreactors were imaged at the confocal microscope in untreated observation chambers consisting of a 24 mm x 60 mm glass cover slide and a 16 mm x 16 mm glass cover slide that were separated by two slices of double-sided sticky tape and sealed with two-component glue. Passive droplets: For the preparation of passive droplets, coacervation is induced right before the encapsulation into the microreactors through the addition of pSS as the last component. The microreactors were sealed and imaged in untreated observation chambers.
Dissolution and reinduction: To exchange the oil phase, 30 µL of a solution containing 10 mM precursor and 5 mM pSS in 200 mM MES at pH 5.3 were added to 300 µL of the oil phase containing the fuel in a 1 mL Eppendorf tube. After 2 h a sample was imaged in an observation chamber as a control for the formation of spherical shells. The residual microreactors in the Eppendorf tube were pipetted into 300 µl of perfluorinated oil containing no fuel. A sample of these microreactors was imaged in an observation chamber to confirm the dissolution of active droplets and active shells. Again after 2 h, the residual microreactors in the oil containing no fuel were pipetted into 50 µl of perfluorinated oil containing 0.5 M DIC. A sample of these microreactors was imaged in an observation chamber.

**(N) Estimating the concentration of product and fuel in the microfluidic droplet**. To validate whether the continuously fueled microreactors contain a steady-state level of product and fuel, we chose an indirect quantification method by HPLC for the product, and quantification by NMR for the fuel. We assumed that the diffusion of fuel from the surrounding oil phase into a microfluidic reactor (r= 5-50 µm) is fast and should be comparable to a two-phase system in a glass vial under vigorous stirring. We used perfluorinated oil without surfactant in these experiments to avoid stabilized emulsions.

Product: 250 µL of an aqueous solution containing 10 mM precursor in 200 mM MES at pH 5.3 was added on top of perfluorinated oil (Novec 7500, 1 mL) in an HPLC vial. We started the reaction network by the addition of 0.5 M DIC into the oil phase and vigorously stirring the reaction mixture. To determine the concentration of the product, 10 µL of the aqueous phase was quenched with 20 µL of an aqueous solution of ethylamine (400 mM) every 30 min. The reaction between the amine and the product yields a stable amide that can be quantified (*3, 4*). We correlated the resulting peak integrals of the amide to the product assuming that the absorption of the amide is equal to the absorption of the precursor (Fig. S1).

Fuel and waste in the steady state: 1.25 mL of 10 mM precursor in 200 mM MES at pH 5.3 were added to 5 mL of perfluorinated oil (Novec 7500) containing 0.5 M DIC. The mixture was vigorously stirred and samples of 25 µL were taken and quenched with 25 µL 640 mM borate buffer at pH 10 with 20 vol% $D_2O$ and 80 mM acetonitrile as a reference. The high pH inhibits the direct DIC hydration and the reaction of the precursor with DIC to ensure that the time until the sample is measured is not influencing the measured concentrations of fuel and waste. Samples are measured by $^1$H-NMR spectroscopy. The concentrations of fuel and waste were then calculated from the peak integrals which were compared to the acetonitrile peak reference (Fig. S1). The chemical shifts of the compared signals were $^1$H NMR (400 MHz, $D_2O$): ACN $\delta$ (ppm) = 2.07 (s, 3H, CH3), DIC $\delta$ (ppm) = 1.22-1.23 (d, 12H, CH3), DIU $\delta$ (ppm) = 1.09-1.10 (d, 12H, CH3).

Variation of the fuel concentration in the microreactor: The concentration of fuel in the aqueous phase was measured for different fuel concentrations in the oil phase to determine the partitioning into the aqueous phase. 1 mL of 200 mM MES at pH 5.3 is added to 10 mL of perfluorinated oil (Novec 7500) containing 0.25 – 1.0 M DIC. The mixtures were vortexed for 20 s and after the phases have separated 250 µl of the aqueous phase are added to 250 µl of 640 mM borate buffer at pH 10 with 20 vol% $D_2O$ and 80 mM acetonitrile as a reference. Samples were measured by $^1$H-NMR spectroscopy. The concentrations of fuel were then calculated from the peak integrals which were compared to the acetonitrile peak reference (Fig. S4D). The chemical shifts of the compared signals were $^1$H NMR (400 MHz, $D_2O$): ACN $\delta$ (ppm) = 2.07 (s, 3H, CH3), DIC $\delta$ (ppm) = 1.22-1.23 (d, 12H, CH3).

**(O) Thermodynamic model for the experimental phase diagram**

In the theoretical model, we use an effective ternary mixture where the influence of fuel, waste, and polyanion is accounted for in an implicit manner; details see section (P). The effective components of the ternary mixture are solvent, the precursor $A$, and the product $B$. For these components, equilibrium concentrations inside and outside were determined experimentally; see Table (1-2). To fit the corresponding experimental diagram, we use a Flory-Huggins free energy density given as (9, 10)

$$f(c_A, c_B) = k_B T \left[ \sum_{i=A,B,S} c_i \log\left(r_i c_i\right) + \sum_{ij=AB,AS,BS} \chi_{ij}\, r_i c_i\, r_j c_j \right] , \tag{3}$$

where $c_A$ and $c_B$ denote the the concentrations of components $A$ and $B$, respectively. In Eq. (3), we have combined the molecular volumes $\nu_A$, $\nu_B$, and $\nu_S$ in the ratios introduced the molecular volume ratios $r_i = \nu_i/\nu_S$. The concentration of the solvent follows from the incompressibility of the mixture, $c_S = 1/\nu_S - r_A c_A - r_B c_B$. This free energy density depends on five parameters: the molecular volume rations, $r_A$ and $r_B$, and the interaction parameters, $\chi_{AB}$, $\chi_{AS}$, and $\chi_{AB}$. We determine these five parameters by fitting experimental measurements of different phase equilibria. At phase equilibria, the chemical potentials $\mu_i = \partial f/\partial c_i$ of components $A$ and $B$ and the osmotic pressure $\Pi = -f + \sum_{i=A,B} c_i \mu_i$ are balanced between the phases. For the measurements, the product was stabilised against hydrolysis by mutating the C terminal aspartic acid for an asparagine. This chemical modification yields a peptide that has the same interaction propensities as the product but is stable, i.e., it does not convert to the precursor. Every measured point in the phase diagram gives three constraints

$$\mu_A\left(c_A^{\mathsf{I}}, c_B^{\mathsf{I}}\right) = \mu_A\left(c_A^{\mathsf{II}}, c_B^{\mathsf{II}}\right) , \tag{4}$$

$$\mu_B\left(c_A^{\mathsf{I}}, c_B^{\mathsf{I}}\right) = \mu_B\left(c_A^{\mathsf{II}}, c_B^{\mathsf{II}}\right) , \tag{5}$$

$$\Pi\left(c_A^{\mathsf{I}}, c_B^{\mathsf{I}}\right) = \Pi\left(c_A^{\mathsf{II}}, c_B^{\mathsf{II}}\right) . \tag{6}$$

We obtained the five unknown parameters by simultaneously minimizing the deviations for the resulting 13 conditions. The best fit was obtained for $r_A = 35.1$, $r_B = 19.4$, $\chi_{AB} = -0.18$, $\chi_{AS} = 0.78$, and $\chi_{AS} = 1.29$. In Sup. fig. 12, we show the corresponding phase diagram together with the experimental measured concentration values. The last thermodynamic parameter needed for our model is the surface tension $\gamma$. Since $\gamma$ is difficult to estimate within our experimental setup, we use the value $\gamma = 75\,\mu$N m$^{-1}$ which is thousand times smaller than the air-water interfacial tension. Our value is in good agreement with surface tensions measured for similar coacervates (11). Note that the used value is slightly larger than for biological condensates (12).

**(P) Sharp interface model for the kinetics of active droplets and active spherical shells**

In general, diffusion is driven by spatial gradients of chemical potentials, while reactions minimize the difference in chemical potentials between products and reactants. For the following, we consider linearized kinetic equations, where diffusion is driven by spatial gradients of concentrations and the reaction

$$A \rightleftharpoons B \,, \tag{7}$$

is driven by the differences in concentrations of the respective components. The resulting kinetic parameters in terms of diffusivities and rate coefficients are then determined via experimental measurements, see experimental section. Therefore, the dynamical equation for $c_A$ and $c_B$ read (13, 14)

$$\partial_t c_i^\alpha = D_i{}^\alpha \nabla^2 c_i{}^\alpha + k_i^\alpha c_j{}^\alpha - k_{ji}^\alpha c_i{}^\alpha \,. \tag{8}$$

In the case of spherical active droplets, there are two different domains with $\alpha =$ I,II, where I denotes the dense phase of droplet and II denotes the dilute phase outside the droplet. For spherical symmetric active spherical shells, three different domains exist, $\alpha =$ I,II,III, where I denotes the dilute phase within the core of the spherical shell, III labels the dense phase of the spherical shell, and II correspond to the dilute phase outside the spherical shell.

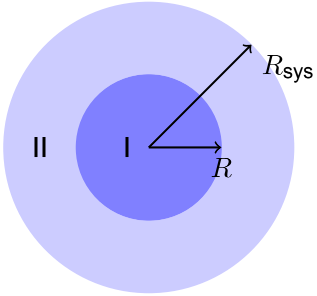


**Sketch I.** Geometry for active droplet.

**(P1) Chemically active droplet**

We calculate the radial symmetric stationary profiles that solve Eq. (8) in the two different domains as a function of the radial coordinate $r$. At the droplet's core ($r = 0$), we impose no flux boundary conditions for the solution of domain I. Similarly, no flux boundary conditions are imposed at system boundary $r = R_{sys}$ for the solution in domain II. The two domains are coupled at the sharp interface at position $r = R$. Here, we impose concentration boundary conditions $c^\alpha_A(R) = a^\alpha$, $c^\alpha_B(R) = b^\alpha$. These concentrations are later determined self-consistently. For these boundary conditions, the stationary solutions for domain I read

$$c^{\text{I}}_A(r) = \frac{b^{\text{I}} - a^{\text{I}}\rho^{\text{I}}_k}{\rho^{\text{I}}_D - \rho^{\text{I}}_k}\frac{R\sinh(r/\lambda^{\text{I}})}{r\sinh(R/\lambda^{\text{I}})} + \frac{a^{\text{I}}\rho^{\text{I}}_D - b^{\text{I}}}{\rho^{\text{I}}_D - \rho^{\text{I}}_k}, \tag{9}$$

$$c^{\text{I}}_B(r) = \rho^{\text{I}}_D\frac{b^{\text{I}} - a^{\text{I}}\rho_k}{\rho^{\text{I}}_D - \rho^{\text{I}}_k}\frac{R\sinh(r/\lambda^{\text{I}})}{r\sinh(R/\lambda^{\text{I}})} + \rho^{\text{I}}_k\frac{a^{\text{I}}\rho^{\text{I}}_D - b^{\text{I}}}{\rho^{\text{I}}_D - \rho^{\text{I}}_k}, \tag{10}$$

while in domain II, the stationary solutions follow

$$c^{\text{II}}_A(r) = \frac{a^{\text{II}}k^{\text{II}}_{BA} - b^{\text{II}}k_{AB}}{k^{\text{II}}_{BA} - k^{\text{II}}_{AB}\rho^{\text{II}}_D}\,\frac{R}{r}\,\frac{\sinh(r/\lambda^{\text{II}}) + \Phi\cosh(r/\lambda^{\text{II}})}{\sinh(R/\lambda^{\text{II}}) + \Phi\cosh(R/\lambda^{\text{II}})} - \frac{a^{\text{II}}k^{\text{II}}_{BA} - b^{\text{II}}k_{AB}\rho^{\text{II}}_D}{k^{\text{II}}_{BA} - k^{\text{II}}_{AB}\rho^{\text{II}}_D}, \tag{11}$$

$$c^{\text{II}}_B(r) = \rho^{\text{II}}_D\frac{a^{\text{II}}k^{\text{II}}_{BA} - b^{\text{II}}k_{AB}}{k^{\text{II}}_{BA} - k^{\text{II}}_{AB}\rho^{\text{II}}_D}\,\frac{R}{r}\,\frac{\sinh(r/\lambda^{\text{II}}) + \Phi\cosh(r/\lambda^{\text{II}})}{\sinh(R/\lambda^{\text{II}}) + \Phi\cosh(R/\lambda^{\text{II}})} - \rho^{\text{II}}_k\frac{a^{\text{II}}k^{\text{II}}_{BA} - b^{\text{II}}k_{AB}\rho^{\text{II}}_D}{k^{\text{II}}_{BA} - k^{\text{II}}_{AB}\rho^{\text{II}}_D}. \tag{12}$$

We have made use of $\lambda^\alpha = \sqrt{D^\alpha_A D^\alpha_B/(D^\alpha_A k^\alpha_{AB} + D^\alpha_B k^\alpha_{BA})}$, $\rho^\alpha_D = -D^\alpha_A/D^\alpha_B$, and $\rho^\alpha_k = k^\alpha_{BA}/k^\alpha_{AB}$, where $\alpha = \text{I}, \text{II}$ indicate the phases.

The coefficient $\Phi = -(\lambda^{\text{II}}R_{\text{sys}}\cosh[\lambda^{\text{II}}R_{\text{sys}}] - \sinh[\lambda^{\text{II}}R_{\text{sys}}])/(\lambda^{\text{II}}R_{\text{sys}}\sinh[\lambda^{\text{II}}R_{\text{sys}}] - \cosh[\lambda^{\text{II}}R_{\text{sys}}])$ ensures the zero flux boundary condition at the system radius.

Finally, we have to determine the four interface concentrations $a^{\text{I}}$, $a^{\text{II}}$, $b^{\text{I}}$, $b^{\text{II}}$ and the position of the interface $R$. For this, we need five constraints. Three of these constraints follow from the assumption of a local equilibrium of phase separation (14)

$$\mu_A(a^{\mathrm{I}}, b^{\mathrm{I}}) = \mu_A(a^{\mathrm{II}}, b^{\mathrm{II}}), \tag{13}$$

$$\mu_B(a^{\mathrm{I}}, b^{\mathrm{I}}) = \mu_B(a^{\mathrm{II}}, b^{\mathrm{II}}), \tag{14}$$

$$\Pi(a^{\mathrm{I}}, b^{\mathrm{I}}) = \Pi(a^{\mathrm{II}}, b^{\mathrm{II}}) - \frac{2\gamma}{R} . \tag{15}$$

Furthermore, the total of $A$ and $B$ have to be conserved in the system,

$$V_{\mathrm{sys}}\, \bar{c}_{\mathrm{tot}} = \frac{4}{3}\pi \left[ (a^{\mathrm{I}} + b^{\mathrm{I}}) R^3 + (a^{\mathrm{II}} + b^{\mathrm{II}})(R_{\mathrm{sys}}^3 - R^3) \right] , \tag{16}$$

where $c_{\mathrm{tot}} = c_A + c_B$ is the conserved quantity associated to the chemical reaction in Eq. (7). Finally, the conservation law at the interface in the stationary state requires

$$j_i^{\mathrm{I}}(R) = j_i^{\mathrm{II}}(R) , \tag{17}$$

where $j_i^\alpha = -D_i^\alpha \partial_r c_i^\alpha$ is radial component of the material flux density. Note that the stationary solution implies that $j_A^\alpha(r) = -j_B^\alpha(r)$. Thus, the constraint Eq. (17) is fulfilled simultaneously for both components $A$ and $B$, making the total number of independent constraints equal to five.

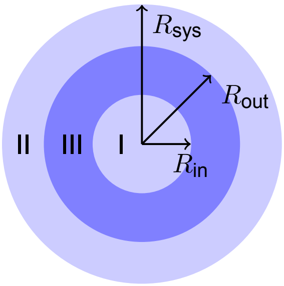


**Sketch II.** Geometry for active spherical shell.

**(P2) Chemically active spherical shell**

In the spherical shell state, the stationary concentration profiles in domain I and II have the same functional form as in the active droplet state. In addition, there is the shell domain III between domain I and domain II. Here, the stationary solutions of Eq. (8), with concentration boundary conditions on the interfaces left and right

$$c_A^{\text{III}}(r) = K_1 \frac{\sinh\big(r/\lambda^{\text{III}}\big)}{r} + K_2 \frac{\cosh\big(r/\lambda^{\text{III}}\big)}{r} + K_3 + \frac{K_4}{r}, \tag{18}$$

$$c_B^{\text{III}}(r) = \rho_D^{\text{III}} \left[ K_1 \frac{\sinh\big(r/\lambda^{\text{III}}\big)}{r} + K_2 \frac{\cosh\big(r/\lambda^{\text{III}}\big)}{r} \right] + \rho_k^{\text{III}} \left[ K_3 + \frac{K_4}{r} \right] . \tag{19}$$

Here, we have abbreviated

$$\begin{aligned} K_1 = \frac{\text{csch}[(R_{\text{out}} - R_{\text{in}})/\lambda^{\text{III}}]}{K_{BA}^{\text{III}} - \rho_D^{\text{III}} K_{AB}^{\text{III}}} & \Big[ (b^{\text{III,in}} K_{AB}^{\text{III}} - a^{\text{III,in}} K_{BA}^{\text{III}}) R_{\text{in}} \cosh\Big(R_{\text{out}}/\lambda^{\text{III}}\Big) \\ & - (b^{\text{III,out}} K_{AB}^{\text{III}} - a^{\text{III,out}} K_{BA}^{\text{III}}) R_{\text{out}} \cosh\Big(R_{\text{in}}/\lambda^{\text{III}}\Big) \Big] , \end{aligned} \tag{20}$$

$$\begin{aligned} K_2 = \frac{\text{csch}[(R_{\text{out}} - R_{\text{in}})/\lambda^{\text{III}}]}{K_{BA}^{\text{III}} - \rho_D^{\text{III}} K_{AB}^{\text{III}}} & \Big[ (b^{\text{III,out}} K_{AB}^{\text{III}} - a^{\text{III,out}} K_{BA}^{\text{III}}) R_{\text{out}} \sinh\Big(R_{\text{in}}/\lambda^{\text{III}}\Big)] \\ & - (b^{\text{III,in}} K_{AB}^{\text{III}} - a^{\text{III,in}} K_{BA}^{\text{III}}) R_{\text{in}} \sinh\Big(R_{\text{out}}/\lambda^{\text{III}}\Big) \Big] , \end{aligned} \tag{21}$$

$$K_3 = \frac{(a^{\text{III,out}} D_A^{\text{III}} + b^{\text{III,out}} D_B^{\text{III}}) R_{\text{out}} - (a^{\text{III,in}} D_A^{\text{III}} + b^{\text{III,in}} D_B^{\text{III}}) R_{\text{in}})}{(D_A^{\text{III}} + D_B^{\text{III}} \rho_k^{\text{III}})(R_{\text{out}} - R_{\text{in}})} , \tag{22}$$

$$K_4 = \frac{(a^{\text{III,in}} - a^{\text{III,out}}) D_A^{\text{III}} + (b^{\text{III,in}} - b^{\text{III,out}}) D_B^{\text{III}}}{(D_A^{\text{III}} + D_B^{\text{III}} \rho_k^{\text{III}})(R_{\text{out}} - R_{\text{in}})} R_{\text{in}} R_{\text{out}} . \tag{23}$$

We introduced $\lambda^{\text{III}} = \sqrt{D_A^{\text{III}} D_B^{\text{III}} / (D_A^{\text{III}} k_{AB}^{\text{III}} + D_B^{\text{III}} k_{BA}^{\text{III}})}$, $\rho_D^{\text{III}} = -D_A^{\text{III}} / D_B^{\text{III}}$, and $\rho_k^{\text{III}} = k_{BA}^{\text{III}} / k_{AB}^{\text{III}}$, in analogy to phases I and II.

Finally, we have to determine the eight interface concentrations $a^{\text{I}}$, $a^{\text{III,in}}$,$a^{\text{III,out}}$, $a^{\text{II}}$, $b^{\text{I}}$, $b^{\text{III,in}}$,$b^{\text{III,out}}$, and $b^{\text{II}}$, and the two positions of the interfaces $R_{\text{in}}$ and $R_{\text{out}}$. For this, we need ten constraints, three of which follow from the assumption of a local equilibrium of phase separation at $R_{\text{in}}$

$$\mu_A(a^{\text{I}}, b^{\text{I}}) = \mu_A(a^{\text{III,in}}, b^{\text{III,in}}) , \tag{24}$$

$$\mu_B(a^{\text{I}}, b^{\text{I}}) = \mu_B(a^{\text{III,in}}, b^{\text{III,in}}) , \tag{25}$$

$$\Pi(a^{\text{I}}, b^{\text{I}}) = \Pi(a^{\text{III,in}}, b^{\text{III,in}}) - \frac{2\gamma}{R_{\text{in}}} , \tag{26}$$

and three from local equilibrium of phase separation at $R_{\text{out}}$

$$\mu_A(a^{\text{III,out}}, b^{\text{III,out}}) = \mu_A(a^{\text{II}}, b^{\text{II}})\,, \tag{27}$$

$$\mu_B(a^{\text{III,out}}, b^{\text{III,out}}) = \mu_B(a^{\text{II}}, b^{\text{II}})\,, \tag{28}$$

$$\Pi(a^{\text{III,out}}, b^{\text{III,out}}) = \Pi(a^{\text{II}}, b^{\text{II}}) - \frac{2\gamma}{R_{\text{out}}}\,. \tag{29}$$

These six equations must be supplemented with a global conservation law

$$\bar{c}_{\text{tot}}\, V_{\text{sys}} = \int dV\,(c_A + c_B)\,, \tag{30}$$

a local conservation law at $R_{\text{in}}$,

$$j_i^{\text{I}}(R_{\text{in}}) = j_i^{\text{II}}(R_{\text{in}})\,, \tag{31}$$

and a local conservation law at $R_{\text{out}}$,

$$j_i^{\text{II}}(R_{\text{out}}) = j_i^{\text{III}}(R_{\text{out}})\,. \tag{32}$$

In contrast to domains I and II, in domain III, $j_A \neq -j_B$, due to the $1/r$ term in the solution of the Laplace equation. Thus, the two flux equations have to be balanced for the components $A$ and $B$ at one interface, respectively. Due to the symmetry of the stationary solutions, one equality is automatically fulfilled at the second interface. Thus, we are left with three independent constraints coming from the local conservation laws at the interfaces Eqs. (31) and (32).

**(Q) Parameter values used in numerical calculations**

If not indicated otherwise, we chose the parameters indicated in Table 6 for the numerical calculations. For the figures shown in this work, we fixed the following parameter values as stated in Table 7.

**(R) Calculations of free energies and free energy rates**

We can estimate the free energy difference between a spherical shell stationary state and the corresponding homogeneous state with the same average concentrations

$$\bar{c}_i = \frac{1}{V_{\text{sys}}} \int dV\, c_i\,, \tag{33}$$

with $i = A, B$. In the following estimates, we will consider spherical shells at steady state corresponding to $R_{\text{sys}} = 35\,\mu$m. We start from the concentrations in the coexisting phases, $c_i^+$ and $c_i^-$, corresponding to the spherical shell average concentration $\bar{c}_i$. Neglecting the interface contribution and considering each phase homogeneous, the free energy in the spherical shell state can be estimated directly via

$$F_{\text{mix}} = V^+ f^+ + (V_{\text{sys}} - V^-) f^-\,, \tag{34}$$

where the free energy density in each phase is $f^{\pm} = f(c_A^{\pm}, c_B^{\pm})$ and $V^+$ is the total volume of the dense phase. With the parameters displayed in Table 6, this difference yields $\text{F}_{\text{mix}} = 2$ nJ.

This free energy can be compared with the total activation free energy, defined as the energy of a $B$ molecule times the number of excess $B$ molecules at the spherical shell steady state

$$F_{\text{act}} = \Delta\omega(\bar{c}_B - c_B^0) V_{\text{sys}}\,, \tag{35}$$

where $c_B^0 = c_{\text{tot}}/[1 + \exp(\Delta\Omega)]$ is the $B$ concentration in the homogeneous equilibrium state, i.e., without fuel present, and $\Delta\Omega$ is the activation free energy of a single $A$ molecule, which is roughly $10\,k_B T$ (15). As outlined in the main text, making use of the parameters in Table 6, we estimate $F_{\text{act}} \simeq 80$ nJ.

Next, we estimate the number of particles that constantly get converted from the precursor $A$ into the product $B$, per unit time and volume,

$$n_{\text{tot}} = \frac{1}{V_{\text{sys}}} \int dV\, c_A\, k_{BA}\,, \tag{36}$$

which determines the total free energy turned over per time and volume

$$J_{\text{tot}} = \Delta\Omega\, n_{\text{tot}}\,. \tag{37}$$

Here, $\Delta\Omega$ denotes the activation energy supplied by fuel to the precursor to form a product. Using the parameters shown in Table 6, $c_{\text{F}} = 8.6$ mM, and $\Delta\Omega \simeq 10k_BT$, we obtain $n_{\text{tot}} = 4 \cdot 10^6\,\text{s}^{-1}\mu\text{m}^{-1}$ and $J_{\text{tot}} = 0.25$ W/L as outlined in the main text.

We can also estimate the free energy flux through the spherical shell interface. Specifically, we calculate the flux of product $B$ through the interface where each product is activated by $\Delta\Omega$

$$J_{\text{int}} = \left[4\pi R_{\text{out}}^2\, j_A^{\text{II}}(R_{\text{out}}) - 4\pi R_{\text{in}}^2\, j_A^{\text{I}}(R_{\text{in}})\right] \Delta\omega / V_{\text{sys}}\,. \tag{38}$$

Using the parameters shown in Table 6, and $\Delta\omega \sim 10\, k_BT$, this leads to an energy influx per unit time of $J_{\text{int}} = 0.198$ W/L. Comparing it to the total power $J_{\text{tot}}$, this leads to an efficiency $J_{\text{int}}/J_{\text{tot}}$ around 70%. In Sup. fig. 16, we explored how $J_{\text{tot}}$ and $J_{\text{shell}}$ vary upon changes in the activation rate, induced by changes of the fuel concentration. Note that in our minimal model, the fuel enters implicitly in the activation rate. As expected, $J_{\text{tot}}$, which represents the total power needed to activate precursor $A$ to product $B$ to their respective values at the non-equilibrium steady state (see definition in Eq. (37)), varies linearly with the fuel concentration. On the other hand, the free energy transported to the spherical shell interface, $J_{\text{int}}$, scales sublinearly with fuel concentration. This is because the higher the fuel the more activation occurs inside, making the $B$ particle flux at the interface less pronounced. As a consequence, the efficiency decreases as fuel concentration increases.

### II. Supplementary Text

**Supplementary discussion 1.** We quantified the interaction strength between the peptides (precursor and product) with pSS by means of ITC. We used Ac-F(RG)$_3$N-NH$_2$ (product*) as a measure for the product (anhydride of precursor) since both molecules are structurally very similar and contain the same positive net charge (*2*).

**Supplementary discussion 2.** For the mechanism evaluation, we wanted to quantify the precursor, product and fuel partitioning in the active droplets. However, for the setup of continuously fueled droplets, a centrifugation assay is not directly possible. Determining the partitioning of the molecules in droplets fueled with a batch of fuel is possible for a specific time point with a centrifugation assay. Still, the molecular composition of the dilute and the dense phase is time-dependent (with respect to fuel addition) and thus this data cannot be used to build a phase diagram. Furthermore, a proceeding of the reactions cannot be prevented during the preparation of these assays. To avoid this time dependence, we prepared passive droplets containing only one of the reactive components at a time. For the precursor and the product, passive droplets at different ratios of precursor and product* (without fuel) were used to measure active droplets in a steady state. To determine the partitioning of the fuel, we prepared droplets consisting only of product* and pSS (no precursor) which were therefore passive as well.

### III. Supplementary tables

| Total concentrations | | | Precursor | | | |
|---|---|---|---|---|---|---|
| Precursor [mM] | Product* [mM] | pSS [mM] | $c_{in}$ [mM] | $c_{out}$ [mM] | $V_{coa}/V_{dilute}$ [vol%] | $K_{partitioning}$ [$c_{in}/c_{out}$] |
| 20 | 0 | 5 | 309 ± 323 | 18.2 ± 1.5 (91%) | 0.57 ± 0.10 | 17 ± 19 |
| 9.5 | 0.5 | 5 | 782 ± 584 | 7.8 ± 0.9 (82%) | 0.22 ± 0.04 | 101 ± 88 |
| 9 | 1 | 5 | 734 ± 229 | 6.1 ± 0.5 (67%) | 0.40 ± 0.05 | 121 ± 49 |
| 8 | 2 | 5 | 655 ± 236 | 6.2 ± 0.3 (78%) | 0.28 ± 0.05 | 106 ± 43 |
| 0 | 2 | 5 | - | - | - | - |

***Table 1:*** *Experimentally determined values for the partitioning of the precursor in passive droplets for the phase diagram (sec. I.(I)).*

| Total concentrations | | | Product* | | | |
|---|---|---|---|---|---|---|
| Precursor [mM] | Product* [mM] | pSS [mM] | $c_{in}$ [mM] | $c_{out}$ [mM] | $V_{coa}/V_{dilute}$ [vol%] | $K_{partitioning}$ [$c_{in}/c_{out}$] |
| 20 | 0 | 5 | - | - | - | - |
| 9.5 | 0.5 | 5 | 176 ± 48 | 0.11 ± 0.03 (23%) | 0.22 ± 0.04 | 1545 ± 904 |
| 9 | 1 | 5 | 233 ± 35 | 0.07 ± 0.02 (7%) | 0.40 ± 0.05 | 3360 ± 1645 |
| 8 | 2 | 5 | 692 ± 139 | 0.10 ± 0.0 (5%) | 0.28 ± 0.05 | 7146 ± 3258 |
| 0 | 2 | 5 | 699 ± 434 | 0.78 ± 0.08 (39%) | 0.18 ± 0.05 | 901 ± 653 |

***Table 2:*** *Experimentally determined values for the partitioning of the product* in passive droplets for the phase diagram (sec. I.(I)).*

| Total concentrations | | | Fuel | | | |
|---|---|---|---|---|---|---|
| Precursor [mM] | Product* [mM] | pSS [mM] | $c_{in}$ [mM] | $c_{out}$ [mM] | $V_{coa}/V_{dilute}$ [vol%] | $K_{partitioning}$ [$c_{in}/c_{out}$] |
| 0 | 2 | 5 | 14.4 ± 17.3 | 9.98 (99.8%) | 0.10 ± 0.01 | 1.44 ± 1.73 |

***Table 3:*** *Experimentally determined values for the partitioning of the fuel in passive droplets (sec. I.(I)).*

| Diffusivity | Spherical shell | Droplet |
|---|---|---|
| $D_{precursor}$ [µm$^2$/s] | 0.060 ± 0.010 | 0.042 ± 0.003 |
| $D_{product*}$ [µm$^2$/s] | 0.029 ± 0.008 | 0.032 ± 0.003 |

***Table 4:*** *Experimentally determined values for the diffusivities in the active droplet and the active spherical shell (sec. I.(L), Fig.S8).*

| reaction constant | Value |
|---|---|
| $k_0$ | $2.9 \cdot 10^{-4}$ s$^{-1}$ |
| $k_1$ | $1.7 \pm 0.08 \cdot 10^{-1}$ M$^{-1}$ s$^{-1}$ |
| K | 1.34 ± 1.0 |
| $k_2$ | $1.2 \pm 0.9 \cdot 10^{-2}$ s$^{-1}$ |

***Table 5:*** *Kinetic parameters determined by the kinetic model (sec. I.(G), Fig. S1).*

| Quantity | Symbol | Value | Reference |
|---|---|---|---|
| Interaction parameter precursor - product | $\chi_{AB}$ | -0.18 | SI II (A) |
| Interaction parameter precursor - solvent | $\chi_{AS}$ | 0.78 | SI II (A) |
| Interaction parameter product - solvent | $\chi_{BS}$ | 1.28 | SI II (A) |
| Precursor relative molecular volume | $r_A$ | 35.1 | SI II (A) |
| Product relative molecular volume | $r_B$ | 19.4 | SI II (A) |
| Surface tension | $\gamma$ | $75\,\mu$N m$^{-1}$ | SI IV (A) |
| Activation rate outside the drop | $k^{\mathrm{II}}_{BA}$ | 0.17 $c_{\mathrm{F}}$ M$^{-1}$s$^{-1}$ | SI I (G) and Tab.5 |
| Activation rate inside the drop | $k^{\mathrm{I}}_{BA}$ | 0.51 $c_{\mathrm{F}}$ M$^{-1}$s$^{-1}$ | SI I (G) and Tab.5 |
| Deactivation rate outside the drop | $k^{\mathrm{II}}_{AB}$ | 0.012 s$^{-1}$ | SI I (G) and Tab.5 |
| Deactivation rate inside the drop | $k^{\mathrm{I}}_{AB}$ | 0.012 s$^{-1}$ | SI I (G) and Tab.5 |
| Diffusion coefficient of $A$ outside the drop | $D^{\mathrm{II}}_{A}$ | 300 $\mu$m$^2$ s$^{-1}$ | SI I (N) and Tab.4 |
| Diffusion coefficient of $B$ outside the drop | $D^{\mathrm{II}}_{B}$ | 300 $\mu$m$^2$ s$^{-1}$ | SI I (N) and Tab.4 |
| Diffusion coefficient of $A$ inside the drop | $D^{\mathrm{I}}_{A}$ | 0.04 $\mu$m$^2$ s$^{-1}$ | SI I (N) and Tab.4 |
| Diffusion coefficient of $B$ inside the drop | $D^{\mathrm{I}}_{B}$ | 0.0073 $\mu$m$^2$ s$^{-1}$ | SI I (N) and Tab.4 |
| Activation rate outside the spherical shell | $k^{\mathrm{II}}_{BA}$ | 0.17 $c_{\mathrm{F}}$ M$^{-1}$s$^{-1}$ | SI I (G) and Tab.5 |
| Activation rate in the spherical shell shell | $k^{\mathrm{III}}_{BA}$ | 0.51 $c_{\mathrm{F}}$ M$^{-1}$s$^{-1}$ | SI I (G) and Tab.5 |
| Activation rate in the spherical shell core | $k^{\mathrm{I}}_{BA}$ | 0.17 $c_{\mathrm{F}}$ M$^{-1}$s$^{-1}$ | SI I (G) and Tab.5 |
| Deactivation rate outside the spherical shell | $k^{\mathrm{II}}_{AB}$ | 0.012 s$^{-1}$ | SI I (G) and Tab.5 |
| Deactivation rate in the spherical shell shell | $k^{\mathrm{III}}_{AB}$ | 0.012 s$^{-1}$ | SI I (G) and Tab.5 |
| Deactivation rate in the spherical shell core | $k^{\mathrm{I}}_{AB}$ | 0.012 s$^{-1}$ | SI I (G) and Tab.5 |
| Diffusion coefficient of $A$ outside the spherical shell | $D^{\mathrm{II}}_{A}$ | 300 $\mu$m$^2$ s$^{-1}$ | SI I (N) and Tab.4 |
| Diffusion coefficient of $B$ outside the drop | $D^{\mathrm{II}}_{B}$ | 300 $\mu$m$^2$ s$^{-1}$ | SI I (N) and Tab.4 |
| Diffusion coefficient of $A$ in the spherical shell shell | $D^{\mathrm{III}}_{A}$ | 0.04 $\mu$m$^2$ s$^{-1}$ | SI I (N) and Tab.4 |
| Diffusion coefficient of $B$ inside spherical shell shell | $D^{\mathrm{III}}_{B}$ | 0.0073 $\mu$m$^2$ s$^{-1}$ | SI I (N) and Tab.4 |
| Diffusion coefficient of $A$ in the spherical shell core | $D^{\mathrm{I}}_{A}$ | 300 $\mu$m$^2$ s$^{-1}$ | SI I (N) and Tab.4 |
| Diffusion coefficient of $B$ inside the spherical shell core | $D^{\mathrm{I}}_{B}$ | 300 $\mu$m$^2$ s$^{-1}$ | SI I (N) and Tab.4 |

***Table 6:*** Table with input parameters used in the numerical calculations.

| Figure | Quantity | Symbol | Value |
|---|---|---|---|
| Fig. 3 C and D | System radius | $R_{\text{sys}}$ | 35 $\mu$m |
| Fig. 3 C and D | Fuel concentration | $c_{\text{fuel}}$ | 8.6 mM |
| Fig. 3 E | System radius | $R_{\text{sys}}$ | 17.5 $\mu$m |
| Fig. 3 E | Fuel concentration | $c_{\text{fuel}}$ | 8.6 mM |
| Fig. 3 G | System radius | $R_{\text{sys}}$ | 25 $\mu$m |
| Fig. 3 G | Fuel concentration | $c_{\text{fuel}}$ | 10 mM |
| Fig. 3 H | System radius | $R_{\text{sys}}$ | 25 $\mu$m |
| Fig. 3 H | Fuel concentration | $c_{\text{fuel}}$ | 10 mM |
| Fig. 3 H | Dif. coef. of $A$ in inside the shell | $D_A^{\text{III}}$ | as indicated by $D_{\text{Precursor}}$ |
| Fig. 3 H | Dif. coef. of $B$ in inside the shell | $D_B^{\text{III}}$ | $D_A^{\text{III}}/5.5$ |
| Sup. fig. 13 A-D | System radius | $R_{\text{sys}}$ | 20 $\mu$m |
| Sup. fig. 13 A-D | Fuel concentration | $c_{\text{fuel}}$ | 10 mM |
| Sup. fig. 13 B | Ratio of dif. inside the drople | $\frac{D_A^{\text{II}}}{D_A^{\text{II}}}$ | $\{1, 2, 4, 5.5, 10\}$ (ind. by arrow) |
| Sup. fig. 13 C | Surface tension | $\gamma$ | $\{25\mu\text{N}, 75\mu\text{N}, 175\mu\text{N}\}$ (ind. by arrow) |
| Sup. fig. 13 D | Ratio of act. rates | $\frac{k_{\text{BA}}^{\text{I}}}{k_{\text{BA}}^{\text{I}}}$ | $\{0.1, 3, 10\}$ (ind. by arrow) |
| Sup. fig. 14 | System radius | $R_{\text{sys}}$ | 35 $\mu$m |
| Sup. fig. 15 | System radius | $R_{\text{sys}}$ | 25 $\mu$m |
| Sup. fig. 16 | Fuel concentration | $c_{\text{fuel}}$ | 8.6 mM |

***Table 7:*** Table with input parameters used for specific figures.

### IV. Supporting Figures

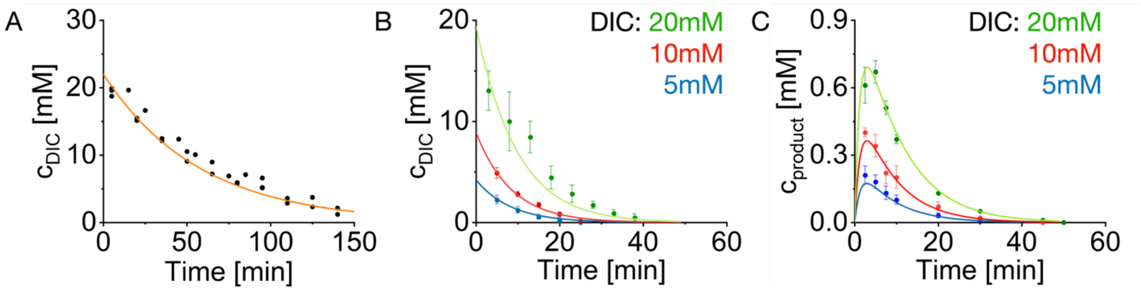


***Supporting figure 1. Determination of the reaction constants. A.*** *Kinetics of the hydration of 20 mM DIC in 200 mM MES buffer at pH 5.3. The DIC concentration was determined by NMR.* ***B.*** *Kinetics of the DIC hydration catalyzed by the precursor. The DIC concentration was determined by NMR. Conditions were 10 mM precursor and 200 mM MES at pH 5.3. Error bars represent the standard deviation of triplicate measurements (N=3). Solid lines represent the fits of the kinetic model.* ***C.*** *Formation of the product upon addition of DIC. The product concentration was determined by analytical HPLC. Conditions were 10 mM precursor and 200 mM MES at pH 5.3. Error bars represent the standard deviation of triplicate measurements (N=3). Solid lines represent the fits of the kinetic model.*

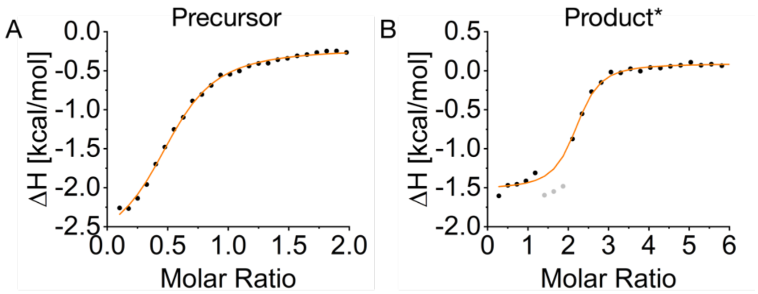


C

| | Precursor | Product* |
|---|---|---|
| $K_D$ / μM | 104.5 ± 11.4 | 2.4 ± 1.2 |
| ΔG / kcal mol$^{-1}$ | -5.4 ± 0.1 | -7.7 ± 0.3 |

***Supporting figure 2. Binding affinity of precursor and product* to pSS. A-B.*** *The change in enthalpy measured by ITC for the interaction between precursor (**A**) or product* (**B**) and pSS. Gray data points were not included in the fitting because they showed signs of coacervation. The solid lines represent the fit of the PEAQ-ITC Analysis software.* ***C.*** *Dissociation constant $K_D$ and free energy ΔG of the interaction between precursor/product* and pSS.*

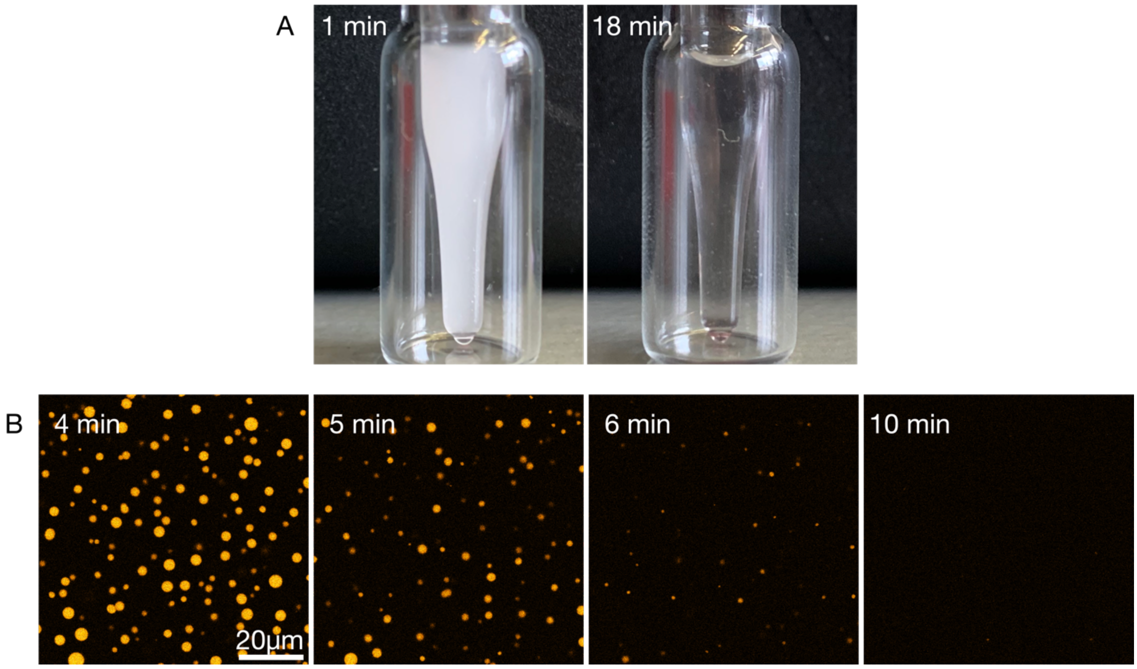


***Supporting figure 3. Batch fueling of active droplets. A.*** *Images of an HPLC inlet containing a solution of 22 mM precursor, 12 mM pSS, and 0.1 µM sulforhodamine B in 200 mM MES buffer at pH 5.3 fueled with 20 mM DIC. After addition of the fuel, the sample turned turbid (1 min) and a clear solution was obtained again after 18 min.* ***B.*** *Confocal microscopy of a solution from A with 0.1 µM sulforhodamine B and fueled with 5 mM DIC. The active droplets grew and fused initially but their size stayed below the size of the active droplets that were fueled with 20 mM DIC (Fig. 1B,C). After around 4 min, they started to dissolve without the formation of spherical shells. Imaging was done in PVA-coated ibidi chambers.*

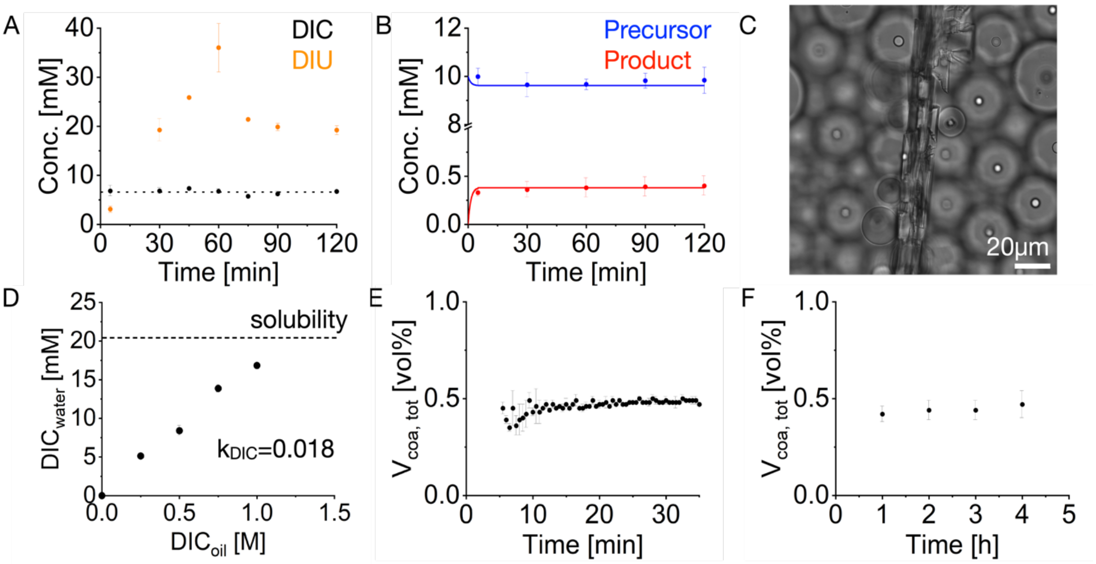


***Supporting figure 4. Total coacervate volumes and reactant concentrations in a steady state.*** ***A.*** *DIC (fuel) and DIU (waste) concentrations under steady-state conditions were determined by NMR. Conditions were 10 mM precursor and 200 mM MES at pH 5.3. To achieve a steady state, perfluorinated oil containing 0.5 M DIC was added on top of the aqueous phase. The dotted line represents the average DIC concentration of 6.6 mM. Error bars represent the standard deviation of triplicate measurements (N=3).* ***B.*** *Average precursor and product concentrations under steady-state conditions were determined by analytical HPLC. Conditions were 10 mM precursor and 200 mM MES at pH 5.3. To achieve a steady state, perfluorinated oil containing 0.5 M DIC was added on top of the aqueous phase. Error bars represent the standard deviation of triplicate measurements (N=3). Solid lines represent the steady-state concentration of DIC calculated by the kinetic model (see methods I.**F** and I.**G**) of 6.6 mM.* ***C.*** *A representative bright field image of a DIU crystal that formed in the oil phase during a steady-state experiment.* ***D.*** *The dependency of the DIC concentration in the oil phase on the DIC concentration in the aqueous phase. The aqueous phase consisted of 200 mM MES at pH 5.3 and the oil phase of Novec 7500 perfluorinated oil. The dotted line represents the DIC concentration in the aqueous phase if an excess of pure DIC was added on top of the aqueous phase without the oil phase. The average partitioning coefficient of DIC between the aqueous phase and the oil phase was $k_{DIC}$ = 0.018. Error bars represent the standard deviation of triplicate measurements (N=3).* ***E.*** *Total coacervate volume in one microfluidic reactor in the first 30 min after the addition of fuel. Conditions were 10 mM precursor, 5 mM pSS, and 200 mM MES at pH 5.3. To achieve a steady state, 0.5 M DIC was added to the oil phase.* ***F.*** *Average total coacervate volume of different microfluidic reactors over the course of 4 h. Conditions were 10 mM precursor, 5 mM pSS, and 200 mM MES at pH 5.3. To achieve a steady state, 0.5 M DIC was added to the oil phase. Error bars represent the standard deviation of at least 30 measurements spread over 3 independent experiments.*

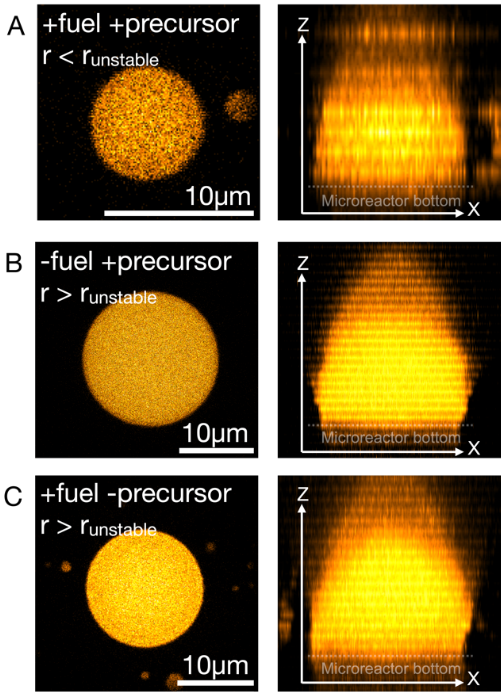


***Supporting figure 5. Wetting of active and passive droplets without transitioning into a shell.*** *All experiments were performed in microfluidic reactors. The left images show the XY-plane at the center of the droplet to demonstrate that no spherical shell was formed. The right images show the XZ-projection of the active/passive droplet to demonstrate the wetting at the bottom of the microreactor.* ***A.*** *A representative active droplet with* $r < r_{unstable}$ *is shown that wetted the bottom of the microreactor but did not transition into a shell. Images were acquired 6 h after the addition of fuel. Conditions were 10 mM precursor, 5 mM pSS, 0.1 µM sulforhodamine B, and 200 mM MES at pH 5.3. To achieve a steady state, 0.5 M DIC was added to the oil phase.* ***B.*** *A representative passive droplet after 3 days is shown with* $r > r_{unstable}$ *that wetted the bottom of a microreactor but did not transition into a spherical shell. Passive droplets were induced by the addition of product* and no fuel was added to the oil phase. Conditions were 9 mM precursor, 1 mM product*, 5 mM pSS, 0.1 µM sulforhodamine B, and 200 mM MES at pH 5.3. Additionally to the fluorosurfactant, 1 w% Krytox 157 FSH was added to the oil phase.* ***C.*** *A representative passive droplet after 6 days is shown with* $r > r_{shell}$ *that wetted the bottom of a microreactor but did not transition into a spherical shell. These passive droplets consist only of product* and pSS without the precursor. Since the added fuel cannot react with the product*, these droplets are thus passive. Passive droplets were induced by the addition of product*. Conditions were 0 mM precursor, 2 mM product*, 5 mM pSS, 0.1 µM sulforhodamine B, and 200 mM MES at pH 5.3 with 0.5 M DIC in the oil phase.*

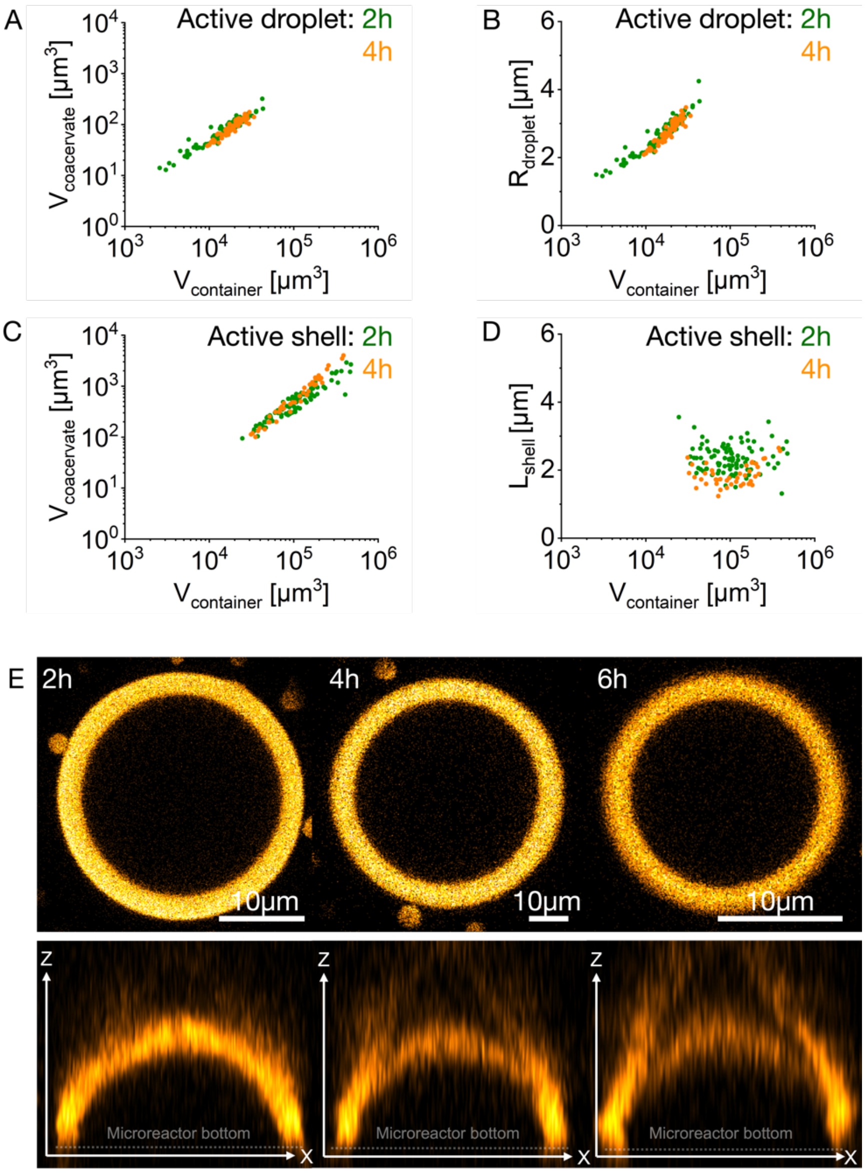


***Supporting figure 6. Stability of active spherical shells.*** *All experiments were performed in microreactors. Conditions were 10 mM precursor, 5 mM pSS, and 200 mM MES at pH 5.3 with 0.1 µM sulforhodamine B. 0.5 M DIC was added to the oil phase to achieve a steady state.* ***A, C.*** *The volume of the total coacervate material 2 h and 4 h after the induction of coacervation is shown for every individual microreactor that contained an active droplet (**A**) or an active shell (**C**). The volume of active droplets and active shells as well as the threshold upon which shells were formed did not change between 2 h and 4 h.* ***B, D.*** *The radius of the active droplet (**B**) as well as the shell thickness $L_{shell}$ of the active shell (**D**) in every individual microreactor is shown 2 h and 4 h after the induction of coacervation. The radius of active droplets and $L_{shell}$ of active shells as well as the threshold upon which shells are formed did not change between 2 h and 4 h. All experiments were performed in triplicate (N=3).* ***E.*** *A representative active shell 2 h, 4 h, and 6 h after the induction of coacervation. In the upper row, the XY-plane of an active shell is shown and in the bottom row, the XZ-plane through the middle of the 3D projection of the respective active shell is shown.*

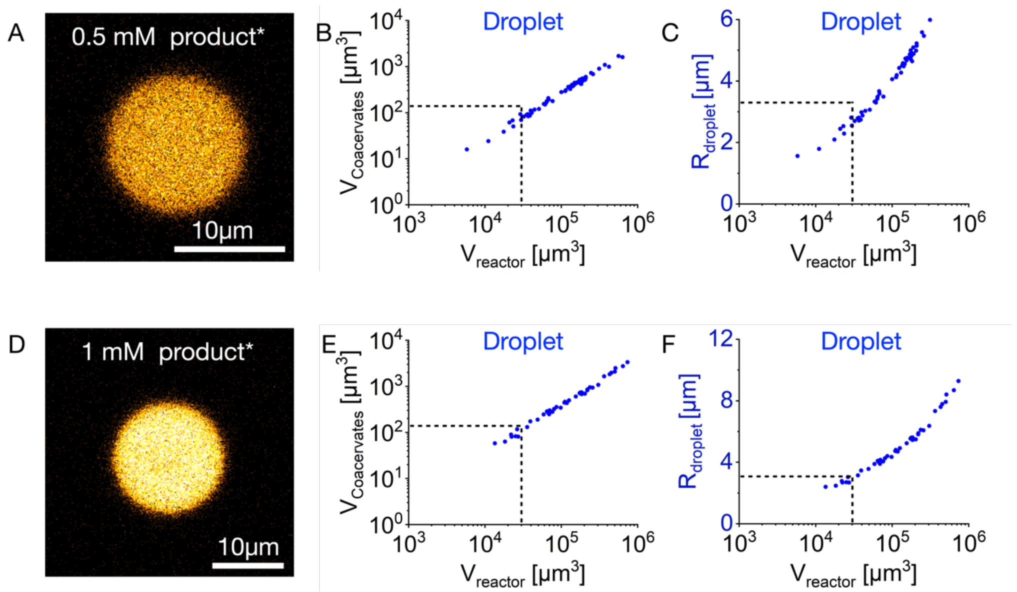


***Supporting figure 7. Passive droplets do not transition into shells.*** *All experiments were performed in microfluidic reactors. For passive droplets, coacervation was induced by the addition of product* instead of the addition of fuel. Images were acquired 4 h after the induction of coacervation. Imaging was done in the same microreactors as for active coacervate-based droplets, but no DIC was added to the oil phase. Dashed lines represent the container size, coacervate volume and size upon which spherical shell formation was observed for active droplets. All experiments were performed in triplicate (N=3).* ***A-C.*** *Conditions were 9.5 mM precursor, 0.5 mM product*,5 mM pSS, 0.1 µM sulforhodamine B, and 200 mM MES at pH 5.3. To show that passive droplets do not transition into shells, a micrograph of a representative passive coacervate-based droplet (**A**), the volume of the total coacervate material (**B**) for every individual microreactor, and the radius (**C**) of the passive droplet in every individual microreactor is shown.* ***D-F.*** *Conditions were 9 mM precursor, 1 mM product*, 5 mM pSS, 0.1 µM sulforhodamine B, and 200 mM MES at pH 5.3. To show that passive droplets do not transition into shells, a micrograph of a representative passive droplet (**D**), the volume of the total coacervate material (**E**) for every individual microreactor, and the radius (**F**) of the passive droplet in every individual microreactor is shown.*

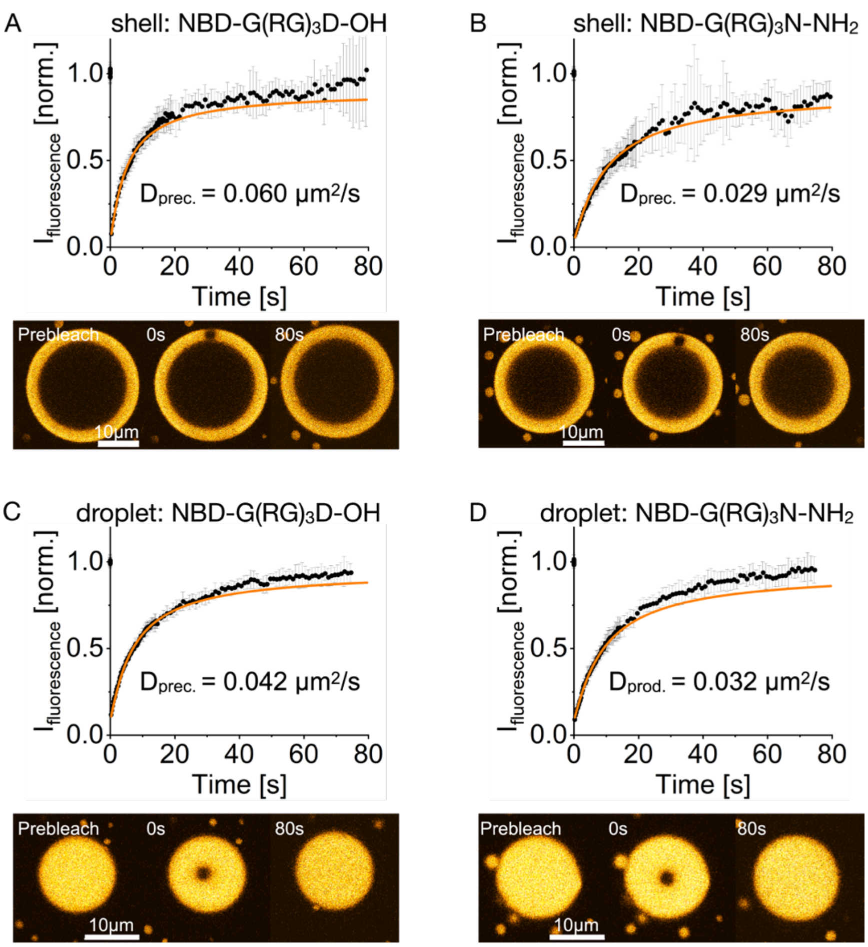


***Supporting figure 8. FRAP of active droplets and active spherical shells.*** *All experiments were performed in microreactors. Conditions were 10 mM precursor, 5 mM pSS, and 200 mM MES at pH 5.3 with 1 µM of the respective dye. To achieve a steady state, 0.5 M DIC was added to the oil phase.* ***A.*** *FRAP of the labeled precursor in active shells 3 h after the induction of coacervation. Error bars represent the standard deviation of 10 different measurements (N=10). The time series shows the fluorescence recovery of a representative active shell.* ***B.*** *FRAP of the labeled product* in active shells 3 h after the induction of coacervation. Error bars represent the standard deviation of 10 different measurements (N=10). The time series shows the fluorescence recovery of a representative active shell.* ***C.*** *FRAP of the labeled precursor in active droplets 1 h after the induction of coacervation. Active droplets were imaged before they transition into a spherical shell. Error bars represent the standard deviation of 10 different measurements (N=10). The time series shows the fluorescence recovery of a representative active droplet.* ***D.*** *FRAP of the labeled product* in active droplets 1 h after the induction of coacervation. Active droplets were imaged before they transition into a spherical shell. Error bars represent the standard deviation of 10 different measurements (N=10). The time series shows the fluorescence recovery of a representative active droplet.*

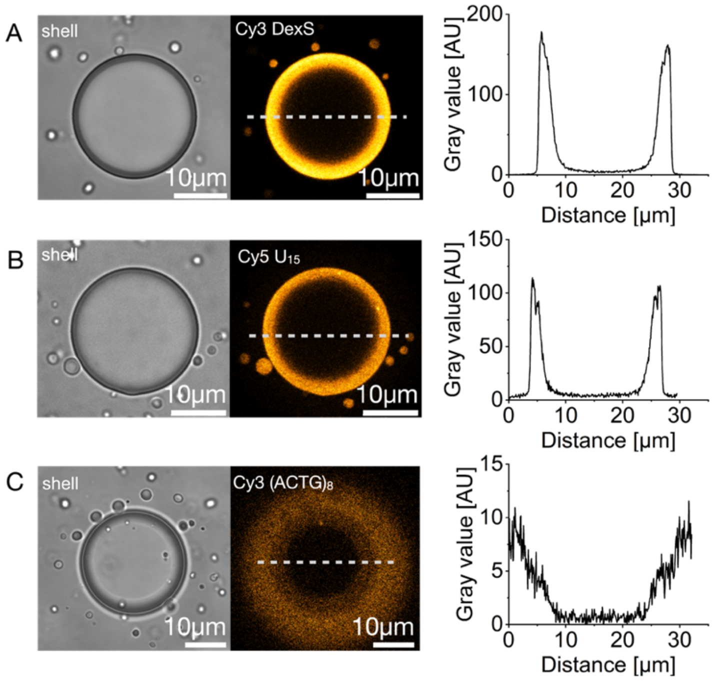


***Supporting figure 9. Partitioning of fluorescent molecules into the active spherical shells.*** *Conditions were 10 mM precursor, 5 mM pSS, and 200 mM MES at pH 5.3 with 1 µM of the respective dye. To achieve a steady state, 0.5 M DIC was added to the oil phase. Active spherical shells were imaged 3 h after the induction of coacervation.* ***A.*** *Bright-field image of an active shell containing Cy3 labeled dextran sulfate (Cy3 DexS, excitation at 552* nm). *The line profile of the fluorescence along the dotted line is shown.* ***B.*** *Bright-field image of an active shell containing Cy5 labeled 15-mer of oligouridylic acid (Cy5 $U_{15}$, excitation at 638* nm). *The line profile of the fluorescence along the dotted line is shown.* ***C.*** *Bright-field image of an active shell containing Cy3 labeled $(ATCG)_8$ (Cy3 $(ATCG)_8$, excitation at 552* nm). *The line profile of the fluorescence along the dotted line is shown.*

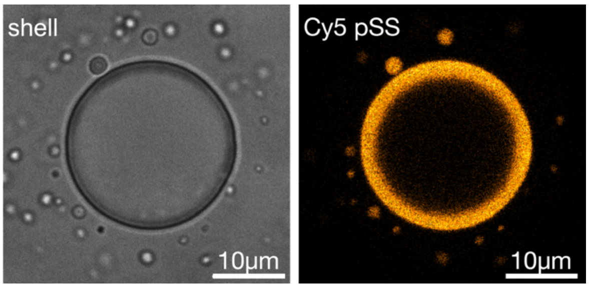


***Supporting figure 10. Partitioning of Cy5-pSS into active spherical shells.*** *Conditions were 10 mM precursor, 5 mM pSS, and 200 mM MES at pH 5.3 with 1 μM Cy5-pSS. To achieve a steady state, 0.5 M DIC was added to the oil phase. The active spherical shell was imaged 2 h after the induction of coacervation. Left: Bright-field image of the active spherical shell. Right: the fluorescence of Cy5-pSS excited at 638* nm.

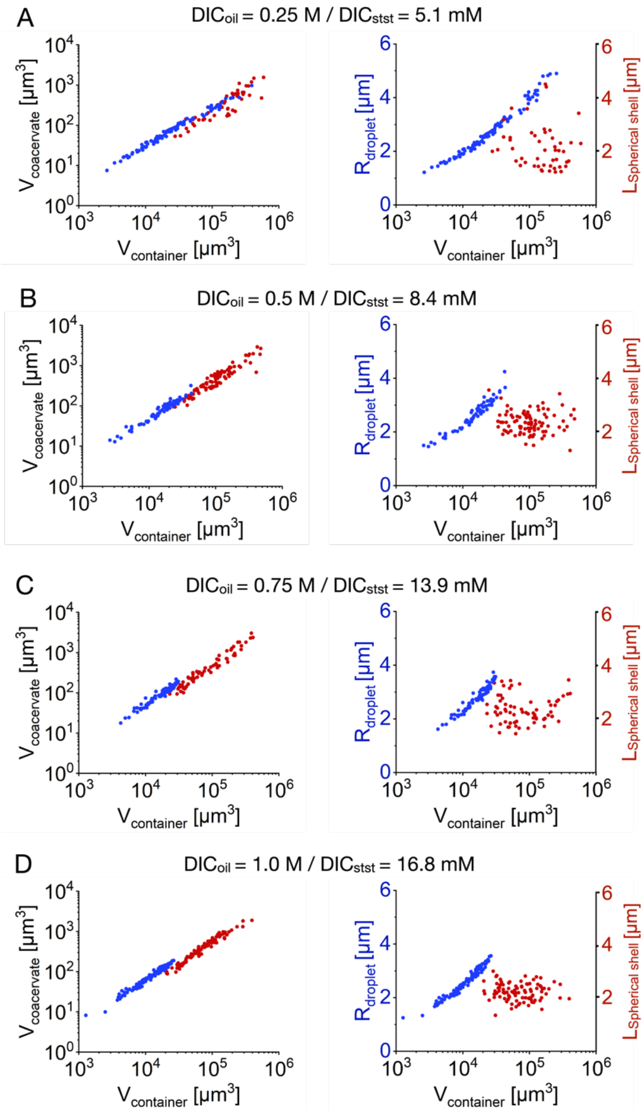


***Supporting figure 11. Spherical shell formation with varying fuel concentrations.*** *Conditions were 10 mM precursor, 5 mM pSS, and 200 mM MES at pH 5.3 with 0.1 µM sulforhodamine B. The indicated amount of $DIC_{oil}$ was added to the oil phase to achieve a steady-state concentration of $DIC_{stst}$ in the microfluidic reactors. Measurements on active droplets and shells were conducted 0.5 h after the spherical shells were formed.* ***A-D.*** *Left: The volume of the total coacervate material is shown for every individual microreactor that contained an active droplet (blue) or an active shell (red). Right: The radius of the active droplet (blue), as well as the shell thickness L (red) of the active shell in every individual microreactor, is shown. All experiments were performed in triplicate (N=3).*

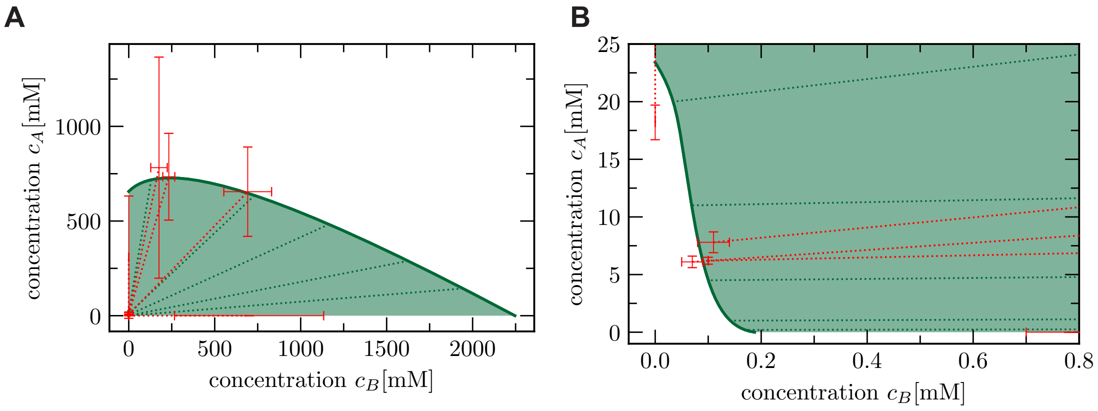


***Supporting figure 12. Phase diagram.*** *Five independent measurements (red data points with error bars) of different phase equilibria were used to fit the binodal line (green solid line) where two phases can coexist. The two corresponding concentration measurements are connected by a red dotted line, while a representative selection of tie lines of the model are shown by the green dotted lines.*

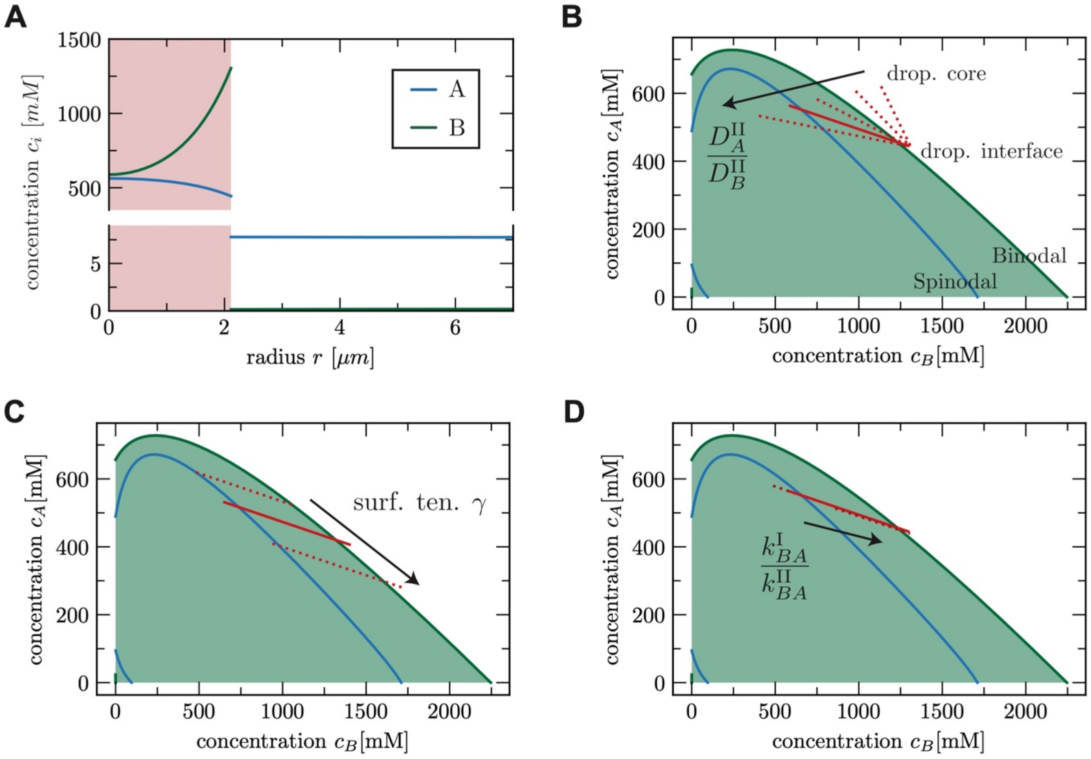


***Supporting figure 13. Influence of kinetic parameters. A** Concentration values of $A$ and $B$ as a function of the radial distance r from the droplet center (red shaded domain is the droplet inside). **B-D** Concentration profiles of the components $A$ and $B$ of the droplet domain (red shaded domain in **A**) in composition space (red line) in addition to the binodal line (green) and the spinodal line (blue). Red solid line corresponds to the kinetic parameters used in the main text (Tab. 6 and 7).*

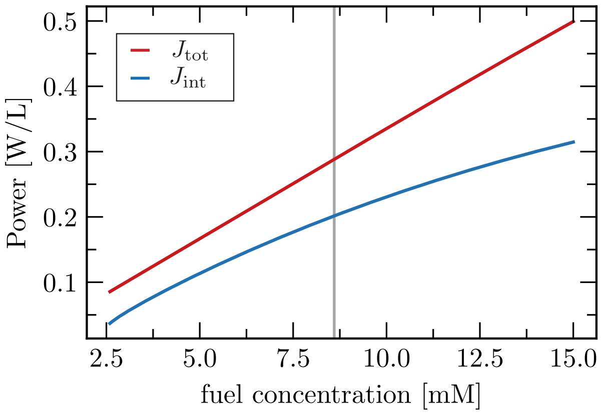


***Supporting figure 14. Influence of fuel concentration on power turn over in the spherical shell state.*** *The red curve represents the power per unit volume needed to constantly activate particles from state $A$ to $B$, $J_{tot}$. We compare it to the free energy transported through the interface corresponding to activated B particles crossing the interface, $J_{int}$ (blue curve). The vertical line corresponds to the fuel concentration value used for the estimates in the main text. In the shown fuel concentration range, the efficiency $J_{int}/J_{tot}$ varies non monotonically between 46% and 70%.*

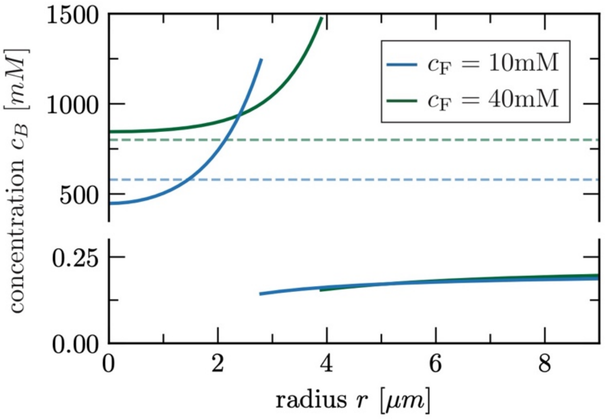


***Supporting figure 15. Influence of fuel concentration on the spinodal instability at the core of the droplet.*** *Radial concentration profiles for two different fuel concentrations are shown. For the lower fuel concentration (blue), the core of a droplet reaches the spinodal concentration (blue dashed). For the higher fuel concentration (green), the stationary droplet is larger. However, the concentration at the core stays above the corresponding spinodal concentration (green dashed) due to weaker gradients resulting from a higher activation inside. For systems with high fuel concentrations, the precursor* $A$ *gets activated and forms the product* $B$ *also inside the droplet.*

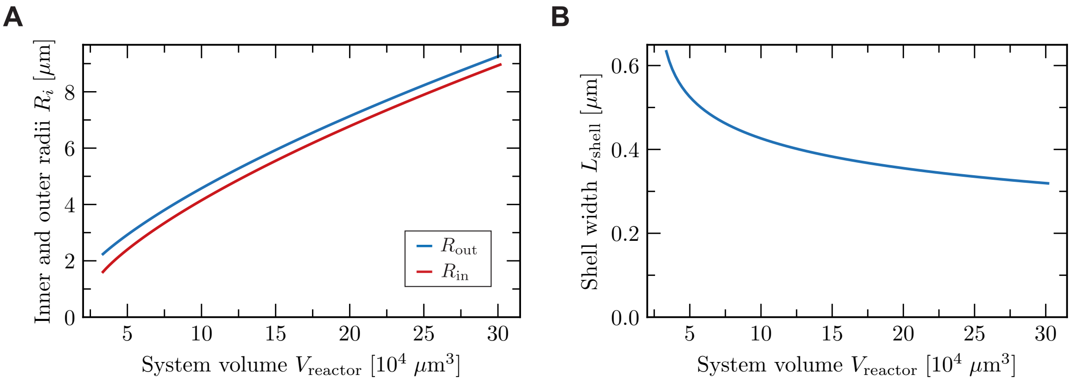


***Supporting figure 16. Influence of the system size on the shell width. A** Both interface radii increase as a function of system volume. **B** Despite the 400 % increase of both radii, the distance between them $L_{shell} = R_{out} - R_{in}$ decreases only by 30 %.*

### V. Supporting Movies

**Supporting movie 1. Spherical shell formation upon dissolution.** *Confocal micrograph timelapse series of a solution of 22 mM precursor, 5 mM pSS in 200 mM MES at pH 5.3 with 0.1 µM sulforhodamine B and fueled with 20 mM DIC. The active droplets grew and fused initially and big droplets formed spherical shells upon dissolution. Imaging was done in PVA-coated ibidi chambers.*

**Supporting movie 2. Dissolution without spherical shell formation.** *Confocal micrograph timelapse series of a solution of 22 mM precursor, 5 mM pSS in 200 mM MES at pH 5.3 with 0.1 µM sulforhodamine B and fueled with 5 mM DIC. The active droplets grew and fused initially but no spherical shells were formed upon dissolution. Imaging was done in PVA-coated ibidi chambers.*

**Supporting movie 3. Formation of a stable active droplet.** *Confocal micrograph timelapse series of a solution of 10 mM precursor, 5 mM pSS in 200 mM MES at pH 5.3 with 0.1 µM sulforhodamine B. To achieve a steady state, 0.5 M DIC was added to the oil phase. The active droplets grew and fused until they reached a stable size of* $r < r_{unstable}$. *The time-lapse series represents the maximum z-projection of a z-stack throughout one microreactor. Imaging was started 10 min after the addition of fuel.*

**Supporting movie 4. Transition of an active droplet into an active spherical shell.** *Confocal micrograph timelapse series of a solution of 10 mM precursor, 5 mM pSS in 200 mM MES at pH 5.3 with 0.1 µM sulforhodamine B. To achieve a steady state, 0.5 M DIC was added to the oil phase. An active droplet with a critical radius larger than* $r_{unstable}$ *transitioned into an active spherical shell. Imaging was done in a microreactor. Imaging was started 1 h after the addition of fuel.*
